# A scalable human-zebrafish xenotransplantation model reveals gastrosome-mediated processing of dying neurons by human microglia

**DOI:** 10.1101/2025.08.27.672578

**Authors:** Ambra Villani, Jana Wittmann, Tamara Wyss, Izaskun Mallona Gonzalez, Irene Santisteban Ortiz, Nathalie Tichy, Monique Pena, Ayush Aditya Pal, Darren Gilmour, Simon T. Schafer, Francesca Peri

## Abstract

Microglia engulf dying neurons through efferocytosis, a critical function in both development and disease. How microglia process the engulfed neuronal material—especially lipids—remains poorly understood, despite its central role in neurodegeneration. Thus, we developed HuZIBRA, a scalable in vivo xenotransplantation model in which human iPSC-derived microglia-like cells (iMGLs) are introduced into the developing zebrafish brain (zf-hiMG), a system characterized by high levels of neuronal cell death and amenable to precise genetic and pharmacological manipulation. We show that human microglia-like cells recognize and engulf apoptotic zebrafish neurons, indicating conserved efferocytic mechanisms. In these cells, engulfed neuronal material accumulates into a distinct, lipid-rich intracellular compartment, the gastrosome, which we also observed in iMGLs placed in a human brain-like environment. The size of the human gastrosome dynamically reflects neuronal cell death levels and is regulated by key genes, including *TREM2* and *SLC37A2*. Pharmacological inhibition of the cholesterol transporter NPC1 induces gastrosome expansion and lipid accumulation, recapitulating pathological features of Niemann-Pick disease type C. Thus, HuZIBRA provides a powerful in vivo platform to uncover cell-autonomous adaptive responses of human microglia to apoptotic and metabolic stress, with the gastrosome emerging as a key integrator of neuronal debris processing and disease-relevant lipid metabolism.

## Introduction

Microglia, the resident immune cells of the brain, play a central role in clearing dead neurons through a process known as efferocytosis. This function is essential during brain development and in several neurodegenerative diseases characterised by high levels of neuronal cell death^1–4^. Interestingly, microglia exhibit similar transcriptional profiles in both contexts, suggesting shared functional responses to neuronal cell death^5,6^ (and reviewed in^7,8^). Efferocytosis of neurons presents unique challenges for microglia, requiring efficient degradation of neuronal components and sorting of resulting byproducts such as lipids. This process becomes especially critical in disease states, where widespread neuronal death drives microglia to adopt an amoeboid morphology often marked by accumulation of lipid inclusions^9–12^. Although the importance of neuronal processing by microglia is increasingly recognized, the underlying cellular mechanisms and the impact of defective processing on microglial shape, plasticity, and function remain largely unknown.

Live imaging studies in zebrafish have shown that microglial phagosomes containing neuronal material fuse with a single compartment called the gastrosome, which serves as a central collection and processing hub^13,14^. This compartment is characterized by a set of unique characteristics such as an electron-lucent lumen containing membrane fragments and lipids^13^. The gastrosome expands in response to increased neuronal cell death and mutations in *slc37a2*^13^. In phagocytic microglia lacking the NPC1 cholesterol transporter, the gastrosome accumulates gangliosides and cholesterol, driving a shift to an amoeboid morphology^14^. This morphological change is further amplified by elevated neuronal cell death, suggesting the gastrosome is an important lipid trafficking compartment highly sensitive to neuronal apoptotic stress.

Given the critical role of microglia in maintaining human brain health, it is essential to determine whether human microglia also possess a gastrosome and how this specialized compartment responds to changes in neuronal cell death. Several studies suggest that genetic differences between animals and humans may lead to variations in microglial behaviour and function^15–17^, indicating that animal models may not fully capture the complexity of human microglia biology. To address this, human-induced pluripotent stem cell-derived microglia-like cells (iMGL) have emerged as a valuable model for studying human-specific mechanisms. However, because the brain microenvironment plays a critical role in shaping microglial identity, responsiveness, and dynamics^18,19^, placing human microglia-like cells in a more physiologically representative context has proven essential. Approaches such as colonizing brain organoids with microglia-like cells^20–23^ (and reviewed in^24,25^) and generating chimeric human/rodent models^17,26–28^ have improved the physiological relevance by providing more accurate brain environments. However, these methods are often time-consuming, resource-intensive, and limited in scalability, which slows experimental throughput and restricts systematic manipulation of microenvironmental factors. Therefore, there is a pressing need to develop scalable approaches that allow precise control over the brain microenvironment, enabling more efficient and detailed investigations into microglial function and their dynamic responses to environmental cues.

In this study, we introduce HuZIBRA (Human Zebrafish Immune-Brain), a scalable in vivo platform for transplanting human iPSC-derived microglia into the zebrafish brain to study their responses to neuronal apoptosis. Live imaging reveals that human microglia complement endogenous microglia and engulf apoptotic neurons, demonstrating high conservation of the efferocytic machinery across species. In these cells we identify the lipid-rich gastrosome, whose size correlates with neuronal death and is regulated by key genes such as *TREM2* and *SLC37A2*. Inhibiting NPC1 expands the gastrosome and induces lipid accumulation, mimicking Niemann–Pick type C pathology. Thus, HuZIBRA provides a physiologically relevant and scalable system to probe neuronal processing and adaptive responses of human microglia in vivo.

## Results

### A scalable *in vivo* pipeline for xenotransplantation of human iPSC-derived microglia-like (iMGL) into the zebrafish embryonic brain

We obtained human iPSC-derived microglia (iMGL) by adopting an established two-step protocol^29^ (see schematics in FIG 1A) where green cytoplasmic (WTC-mEGFP-AAVS1-cl6) or red membrane (WTC-mTagRFPT-CAAX-AAVS1-cl91) fluorescently labelled iPSC were first differentiated into hematopoietic progenitors (iHPC), as confirmed by the expression of typical markers for these cells (SUPPL. FIG1A). Subsequently, iHPC were transferred into microglial differentiation and maturation media (see schematics in FIG 1A). The resulting induced microglia-like cells (iMGLs) displayed a rudimentary branched morphology (FIG 1B), phagocytic activity (FIG 1C) and expressed key microglial markers (FIG 1D and SUPP FIG 1B). Expression of canonical microglial genes in iMGLs was further confirmed both by transcriptome analysis, which showed that these cells cluster together with published human iPSC-derived microglia datasets^29^ (FIG 1E and SUPP FIG 1C, D), and by in situ hybridization (HCR), validating mRNA expression of microglial-specific genes *in vitro* (SUPP FIG 1E). These analyses confirmed that iMGLs were transcriptionally distinct from other cell types, such as iHPC precursors and iPSCs (FIG 1E), and multidimensional scaling revealed strong similarities with both fetal and adult human microglia (FIG 1E).

**Figure 1.**
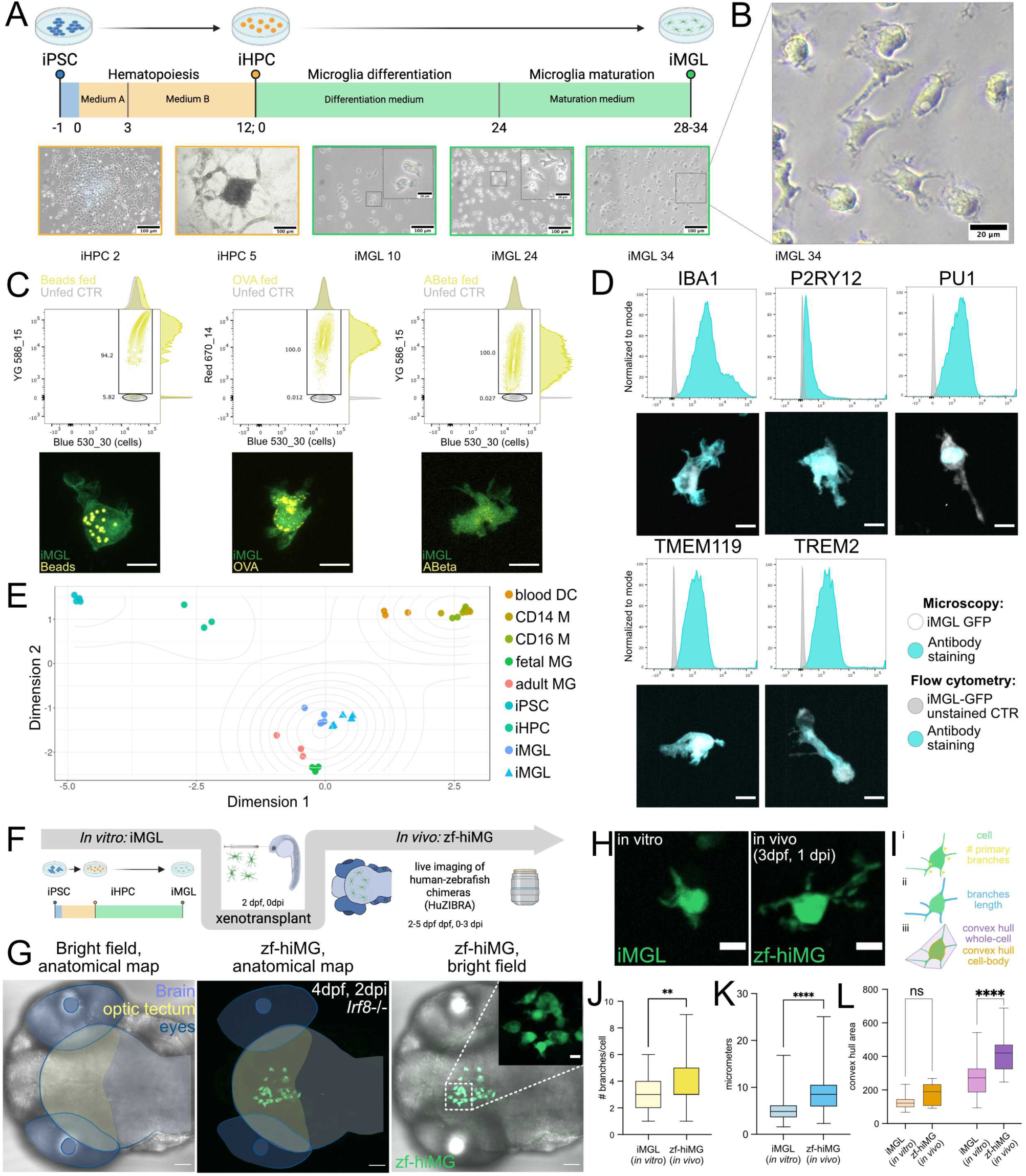
A scalable pipeline for the xenotransplantation of human iPSC-derived microglia (iMGL) into the zebrafish embryonic brain. **(A)** Schematic overview of the two-step differentiation protocol used to generate hiPSC-derived microglia (iMGL) from fluorescently labelled human iPSCs. **(B)** Bright field image of WT iMGL in vitro at day 34 of microglia differentiation; scale bar 20 µm. **(C)** Phagocytic assay in vitro: GFP labelled WT iMGL are fed with fluorescently labelled 1µm latex beads, ovalbumin or Amyloid-beta and analysed by FACS (upper row) or light microscopy (lower row). Scale bars 10 µm. **(D)** Immunostaining of microglial markers on GFP-labelled WT iMGL analysed by FACS and light microscopy; scale bars 10 µm. **(E)** Multidimensional scaling (MDS) comparing WT iMGL obtained in this study (triangles) to iMGL and other cell types obtained by Abud et al. 2019 (circles). **(F)** Schematic of the xenotransplantation protocol of fluorescently labelled iMGL into the embryonic zebrafish optic tectum (zf-hiMG) at 2 days post fertilization (dpf). **(G)** Representative image of a *Irf8^st95^* zebrafish brain at 4dpf, 2 dpi (days post injection), dorsal view, xenotransplanted with GFP-labelled WT iMGL (zf-hiMG); scale bars 50 µm (overview) and 10 µm (zoomed crop). **(H)** Comparison of representative cells *in vitro* (iMGL, left) versus *in vivo* (zf-hiMG, right); scale bars 10 µm. **(I–L)** Comparison of microglial morphology *in vitro* (iMGL) versus *in vivo* (zf-hiMG). Quantification of number of primary branches **(J)** ; branch length (average per cell) **(K)** and convex hull of the cell body or entire cell **(L)** as described in the schematic in **(I)**; N=3, n=81(*in vitro*), n=74 (*in vivo*); N=experiments, n=cells.

To mimic a physiologically relevant brain environment, we transplanted GFP labelled iMGL directly into the optic tectum (OT) of two-day post-fertilization (dpf) zebrafish embryos, a developmental stage marked by extensive neuronal apoptosis and the establishment of the endogenous microglial population (∼25 cells)^30^ (FIG 1F). The combination of high iMGL culture yields and an efficient injection protocol enabled semi-high-throughput transplantation at a rate of approximately 20 chimeric embryos per operator per hour. We first transplanted iMG into *Irf8^st95^* mutants, which lack endogenous microglia and macrophages^31^ (FIG 1G). iMG were detected in the OT of approximately 65% of the transplanted embryos (these cells are hereafter referred to as zf-hiMG; SUPPL. FIG1F, left bar). The number of transplanted zf-hiMG varied, from as few as 1 to over 40 cells per embryo, with an average of 18 (SUPPL. FIG1G, left bar), highlighting the scalability of the model. We confirmed the survival of zf-hiMG (SUPPL. FIG1H) and validated their human microglial identity using *in vivo* Hybridization Chain Reaction analysis (HCR), which demonstrated the expression of key markers, such as *TMEM119*, *P2Y12*, *IBA1*, and *TREM2* (SUPPL. FIG1I). Additionally, TREM2 expression was further confirmed also at the protein level (SUPPL. FIG1J). Together these data provide evidence that zf-hiMG maintain a typical human microglial identity within the zebrafish brain.

The small size and optical transparency of zebrafish embryos enable high spatiotemporal resolution imaging, which we leveraged to investigate the morphology and behavior of transplanted zf-hiMG. To capture these dynamics, we performed whole-brain time-lapse imaging using spinning disk and single-plane illumination microscopy (SPIM)—two high-speed imaging modalities characterized by negligible phototoxicity (Jemielita et al., 2013; Keller and Stelzer, 2008). Transplanted cells exhibited a more elaborate branched morphology compared to iMGLs cultured in vitro (Fig. 1H–J), with significantly longer processes (Fig. 1H–I, K) and enhanced motility, marked by continuous cycles of extension and retraction (Supp. Fig. 1K–L; Fig. 2B; Video 3). Time-lapse imaging further revealed that zf-hiMG were highly dynamic (Fig. 2A; Videos 1–2), suggesting that the zebrafish brain microenvironment supports key microglial properties, including complex morphology and branching dynamics.

**Figure 2.**
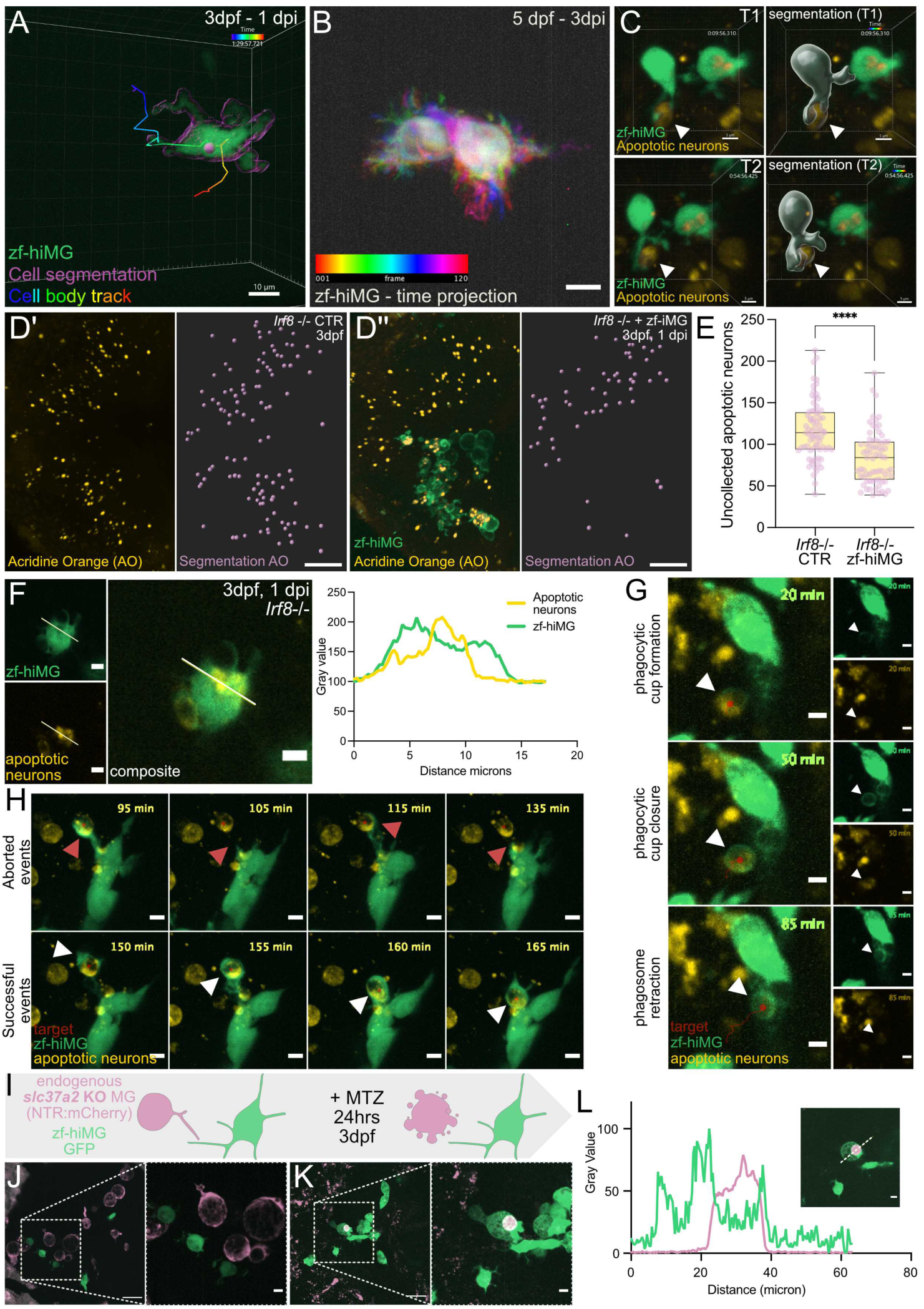
Functional conservation of microglial engulfment mechanisms in a zebrafish–human chimeric model. **(A)** Segmentation and tracking of GFP-labelled WT zf-hiMG cell body in a 3 dpf, 1 dpi *Irf8^st95^* zebrafish OT; 5 minutes time resolution; scale bar 10 µm. **(B)** Time-projection of GFP-labelled WT zf-hiMG in a 5 dpf, 3 dpi zebrafish OT; 1 minute time resolution; scale bar 10 µm. **(C)** Time lapse mages showing transient contacts (arrowheads) between GFP-labelled WT zf-hiMG branches and nearby apoptotic neurons (Tg(nbt:secA5-BFP)) in *Irf8^st95^* 3dpf, 1dpi embryos; 5 min time resolution; scale bar 5 µm. **(D)** Representative images of Acridine Orange (AO) staining and segmentation in 3 dpf *Irf8^st95^* embryos; **(D’)** control and **(D’’)** with membrane RFP labelled WT zf-hiMG transplantation; scale bars 50 µm. **(E)** Quantification of uncollected AO+ apoptotic nuclei in the optic tectum of *Irf8^st95^* embryos ± iMGL transplantation; N=3, n=73 (CTR), n=70 (zf-hiMG). **(F)** Image showing a GFP-labelled WT zf-hiMG transplanted in *Irf8^st95^* containing secA5+ apoptotic neuronal material (Tg(nbt:secA5-BFP)), with quantification of intracellular fluorescent signal; 3 dpf, 1 dpi; scale bar 5 µm. **(G)** Time-lapse imagies showing GFP-labelled WT zf-hiMG transplanted in *Irf8^st95^* and engulfing an apoptotic neuron (Tg(nbt:secA5-BFP)) by forming a phagocytic cup (arrowhead); track of phagosome retraction in red; 5 minutes time resolution; scale bar 5 µm. **(H)** Time-lapse images of a GFP-labelled WT zf-hiMG in *Irf8^st95^* attempting sequential engulfment of a dying neuron (Tg(nbt:secA5-BFP)): top, failed phagocytic cup formation attempt and abortion (red arrowhead); bottom, successful re-engagement and engulfment (white arrowhead); 5 minutes time resolution; scale bars 5 µm. **(I)** Schematic of metronidazole (MTZ)-induced microglial ablation in *slc37a2* KO Tg(fms:Gal4;UAS:nfsB-mCherry) embryos expressing NTR in endogenous zebrafish microglia (MG). **(J–K)** Representative images of **(J)** control and **(K)** MTZ-treated *slc37a2* KO embryos showing selective apoptosis of endogenous NTR+ microglia and uptake of apoptotic material by WT zf-hiMG; scale bars 50 µm (overview) and 10 µm (zoomed crop). **(L)** Quantification of fluorescent apoptotic signal in zf-hiMG 24 hrs after MTZ treatment and correspondent qualitative image; scale bar 5 µm.

To further assess functional engagement with the neural environment, we transplanted iMGL into *Irf8^st95^* mutant zebrafish embryos expressing a fluorescent reporter for apoptotic neurons (Tg(nbt:dLexPR-LexOP:secA5-BFP))^32^. Live imaging revealed transient contacts between zf-hiMG processes and nearby apoptotic neurons (FIG 2C, video 4), suggesting active sensing of the local microenvironment. Next, we asked whether zf-hiMG could survive and function alongside endogenous zebrafish microglia that we labelled with mCherry. Transplantation efficiency was comparable in zebrafish hosts with or without endogenous microglia (SUPP FIG 1F-G), and both populations co-existed within the same brain environment (SUPP FIG 2A and video 5).

Taken together, these findings indicate that this newly developed human-zebrafish xenotransplantation platform (HuZIBRA) represents a powerful tool for *in vivo* exploration of human microglial behaviour and cellular interactions within an accessible brain environment.

### Functional conservation of microglial engulfment mechanisms in a zebrafish-human chimeric model

A key function of microglia during development is the engulfment of dying neurons through efferocytosis^33–36^. In *Irf8^st95^* zebrafish mutant embryos, which lack microglia, uncollected apoptotic neurons accumulate in the developing optic tectum (OT)^31^, as visualized by Acridine Orange (AO) staining (FIG 2D’). Remarkably, transplantation of iMGL into into *Irf8^st95^* embryos significantly reduced the number of uncollected apoptotic nuclei (FIG 2D’-E). This was further validated using a real-time apoptotic reporter (Tg(nbt:dLexPR-LexOP:secA5-BFP))^32^, which revealed fluorescent apoptotic material inside zf-hiMG vesicles, indicating active uptake of labelled apoptotic neurons (FIG 2F). Live time-lapse imaging further revealed that zf-hiMG actively engulf endogenous apoptotic neurons by extending branches and forming phagocytic cups around dying cells (FIG 2G; video 6). These cups either successfully enclosed the target, forming ∼5 µm phagosomes that were retracted toward the microglial cell body, or failed to close, resulting in aborted engulfment attempts (FIG 2H, Video 7), a phenomenon previously observed in zebrafish microglia^32,37^. Notably, failed attempts were often followed by re-engagement and successful engulfment (FIG 2H, video 7).

To further evaluate the efferocytic capacity of zf-hiMG, we tested their response to induced apoptosis in endogenous microglia. To this aim, we used a genetic system allowing selective ablation of zebrafish microglia (Schematic in FIG 2I). GFP-labeled iMGL were transplanted into slc37a2-deficient embryos, in which endogenous microglia expressed a nitroreductase–mCherry fusion protein (Tg(fms:Gal4,UAS:nfsB-mCherry)). These slc37a2-deficient microglia are readily identifiable by their abnormally enlarged gastrosome, a consequence of impaired neuronal digestion^13^ (FIG 2J, SUPP FIG 2B’’, C-C’). Metronidazole (MTZ) treatment triggered Caspase-3-dependent apoptosis specifically in nitroreductase-expressing microglia^30,38–40^ (SUPP FIG 2B, compare B’’ with B’’’), but had no effect in non-transgenic controls^30^ (SUPP FIG 2C-C’). Within 24 hours, red fluorescent apoptotic material was visible inside zf-hiMG (FIG 2K, L), indicating they could recognize and engulf dying endogenous microglia via conserved “find me” and “eat me” signals. Furthermore, zf-hiMG responded to targeted neuronal laser ablation by polarizing and migrating toward these injury sites (SUPP FIG 2D, video 8), indicating their capacity to detect and respond to local damage-associated cues.

Finally, we examined whether zf-hiMG could assist endogenous *slc37a2* zebrafish microglia that have an enlarged gastrosome due to neuronal processing defects ^13^. iMG transplantation into *slc37a2* mutants led to a marked reduction in gastrosome size within the endogenous *slc37a2* microglia population (SUPP FIG 2E-G compare F’’ with E’, quantification in SUPP 2G). Because the size of the gastrosome in these mutants is known to depend on the rate of neuronal engulfment^13^, this reduction suggests that zf-hiMG alleviate the burden by clearing apoptotic neurons, thereby functionally complementing the endogenous microglial population.

Together, these results highlight the evolutionary conservation of efferocytic mechanisms between zebrafish and humans and establish this chimeric model as a powerful platform to study human microglial responses to neuronal cell death in a physiologically relevant context.

### Xenotransplantation of human microglia-like cells into zebrafish reveals gastrosome adaptation under phagocytic stress

Building on our observations that zf-hiMG can recognize apoptotic zebrafish neurons, we next examined how they process this material. Tracking neuron-derived apoptotic cargo within zf-hiMG revealed progressive intracellular aggregation (FIG 3A; Video 9), a phenomenon consistent with the presence of the gastrosome—an electron-lucent phagocytic compartment where engulfed apoptotic material accumulates (Villani et al., 2019). To determine whether a similar process occurs in iMGLs, we exposed these cells in culture to fluorescent latex beads and monitored intracellular cargo distribution over a 24-hour period. The beads progressively clustered within the cells (SUPP FIG 3A), and electron microscopy (EM) revealed the presence of a single enlarged vesicle displaying key features of the gastrosome, including an electron-lucent lumen enclosed by a single membrane and containing membrane fragments^13,14^ (FIG 3B). While no molecular marker uniquely defines the gastrosome, its ultrastructural characteristics, perinuclear position, size enlargement in response to increased neuronal cell death and lipid content, enable its reliable identification across different contexts^13,14,41^. Given that in vitro cultures inherently experience a certain degree of spontaneous apoptosis (Supp. Fig. 3B–C), some human iMGLs also displayed an enlarged single vesicle in the absence of latex beads. As shown by EM and 3D reconstructions, this compartment shares the same morphological features as the gastrosome observed across different microglial contexts^13,14^ (FIG 3C and SUPP FIG 3 D-D’). We next asked whether human microglia that differentiate within a human brain-like environment would also form a gastrosome, pointing to conserved neuronal processing in these cells. Brain organoids provide such a human brain-like environment and neuronal cell death is an intrinsic feature of their growth and maturation^23^. Indeed, we have previously shown that long-term organoid culture leads to necrotic core formation and the emergence of developmentally stressful microenvironments^23^. Consistent with this, EM imaging and 3D segmentation of enlarged amoeboid microglia within organoids revealed a single, perinuclear, electron-lucent vesicle containing cellular debris, further supporting the presence of a gastrosome-like compartment in human microglia under these conditions (FIG 3D-D” and Video 10).

**Figure 3.**
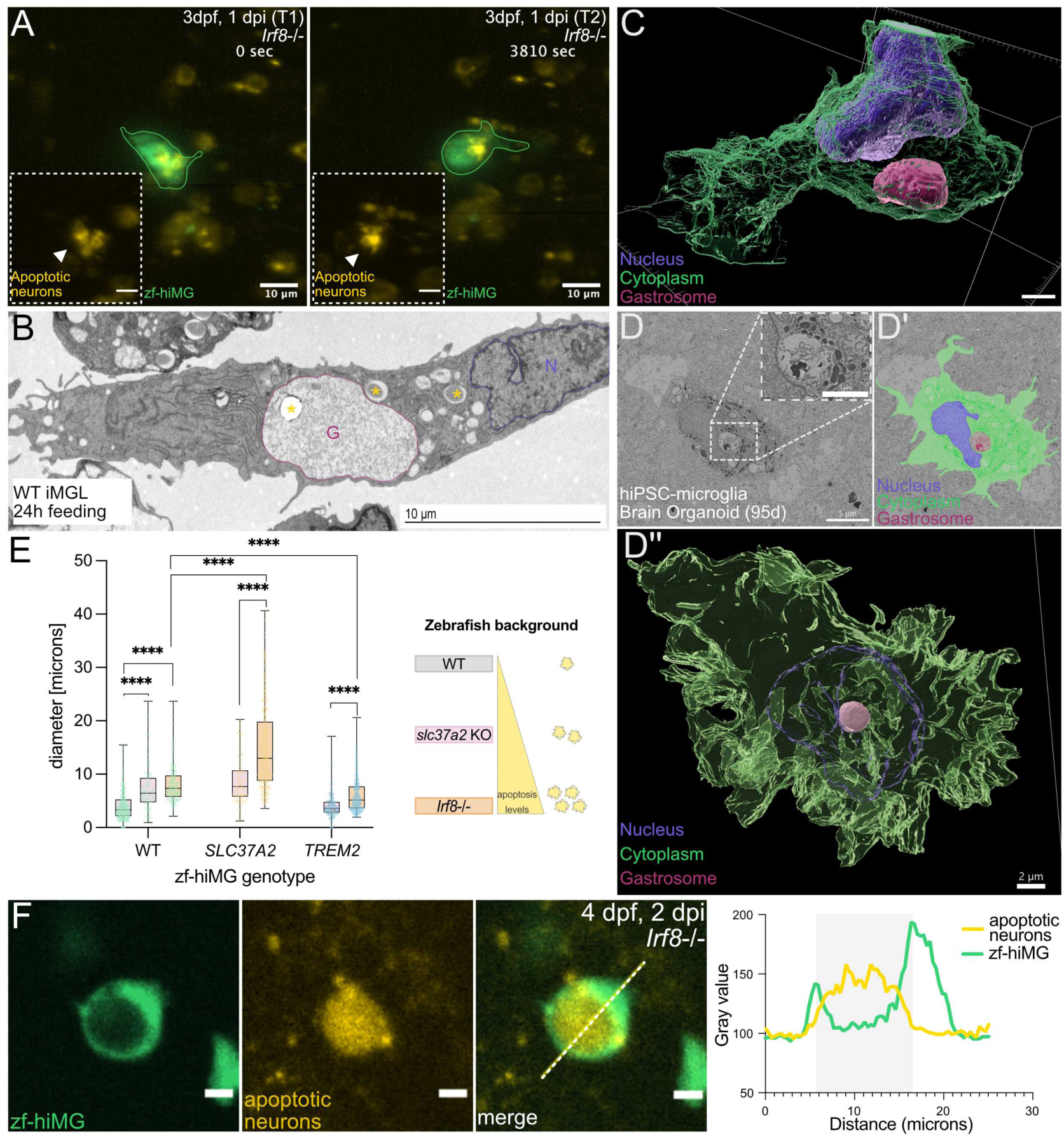
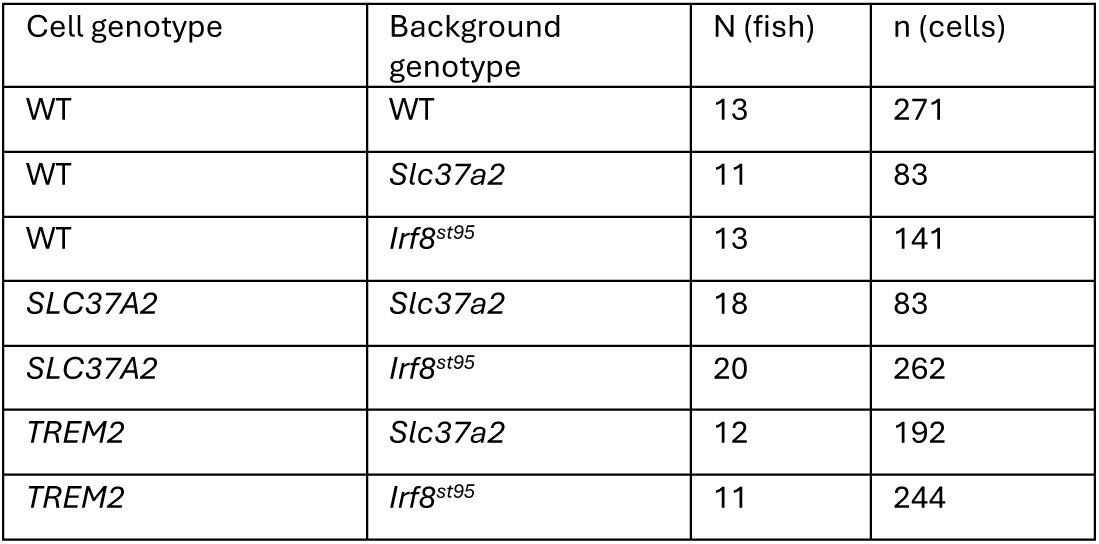
The gastrosome in human microglia-like cells is a conserved, lipid-rich compartment regulated by efferocytic activity. **(A)** Time-lapse images of GFP-labelled WT zf-hiMG in Irf8^st95^ embryos, showing interaction with neurons marked for apoptosis (Tg(nbt:dLexPR-LexOP:secA5-BFP); 30 seconds time resolution; 3 dpf, 1 dpi; scale bars 10 µm (overview) and 5 µm (zoomed crop). **(B)** Electron micrograph (single plane) of 24 hrs bead-fed WT iMGL in monoculture; N = nucleus, G = gastrosome, * = beads-containing phagosomes; scale bar 10 µm. **(C)** 3D volume reconstruction of SEM data showing the cytoplasm (green), nucleus (blue) and gastrosome (purple) in WT iMGL in vitro (monoculture); scale bar 2 µm. **(D)** Single slice from an SEM array of hiPSC-derived microglia integrated into a 95 days old brain organoid; scale bars 5 µm (overview) and 2 µm (zoomed crop); **(D’)** segmentation of **(D)** showing the cytoplasm (green), nucleus (blue) and gastrosome (purple); **(D’’)** 3D volume reconstruction of the segmented cell; scale bar 2 µm. **(E)** Quantification of the gastrosome’s diameter in GFP-labeled zf-hiMG across different zebrafish backgrounds with increasing apoptotic burden: WT (grey), *slc37a2* KO (pink), and *Irf8*⁻/⁻ (orange). Boxplots show WT zf-hiMG (left column), *SLC37A2* zf-hiMG (middle), and *TREM2* zf-hiMG (right); N (fish), n (cells) in the table below (data from 5 independent experiments). **(F)** Representative image and fluorescence intensity quantification of GFP-labelled WT zf-hiMG in *Irf8*-/ background with neuronal apoptotic marker (Tg(nbt:dLexPR-LexOP:secA5-BFP); 4dpf, 2 dpi; scale bar 5 µm.

As high levels of neuronal apoptosis is a hallmark of many human disease contexts, including neurodegenerative disorders and brain injuries, we next examined how the microglial gastrosome responds to apoptotic stress and increased engulfment demands. To this end, we transplanted WT zf-hiMG into three different zebrafish models characterised by different levels of apoptosis: WT (low burden), *slc37a2* mutants (moderate burden)^13^, and *Irf8^st95^* (high burden)^31^. Gastrosome size increased progressively across these conditions, correlating with the apoptotic load (FIG 3E, leftmost column: grey, pink and orange boxplots for WT zf-hiMG). Imaging in the *Irf8^st95^* background, combined with a marker for apoptotic neurons, confirmed that enlarged gastrosomes contained neuron-derived material (FIG 3F). The dose-dependent enlargement underscores the gastrosome’s adaptive responsiveness to increasing levels of apoptotic stress, further supporting its role as a specialized phagocytic compartment for processing engulfed cellular debris.

Given the observed correlation between gastrosome size and neuronal cell death^13,14^, we next asked whether this compartment’s responsiveness would be altered in microglia with impaired phagocytic function. TREM2, a key regulator of microglial phagocytic functions and a well-established risk factor in multiple neurodegenerative diseases^5,42–44^, appeared as a suitable candidate to test this hypothesis. To this end, we generated *TREM2*-deficient hiPSC using CRISPR-Cas9, targeting exon 2 with a previously published sgRNA^44^ (SUPP FIG 4A). We obtained a clone carrying a homozygous small deletion (SUPP FIG 4A’), which led to significantly reduced transcript levels as confirmed by RNA-seq (SUPP FIG 4B). The edited cells retained pluripotency (data not shown) and differentiated efficiently into iMGL (*TREM2* iMGL) expressing canonical microglial markers at levels comparable to WT iMGL (SUPP FIG 4B). Transplantation of *TREM2* zf-hiMG into *Irf8^st95^* embryos resulted in a significantly higher number of uncollected apoptotic neurons compared to embryos transplanted with WT zf-hiMG (FIG 4A). Notably, the number of unengulfed apoptotic neurons was comparable to that observed in non-transplanted *Irf8^st95^* embryos, indicating that *TREM2*-deficient zf-hiMG fail to compensate for the absence of endogenous zebrafish microglia. This functional impairment is consistent with previous studies in other systems^45–50^. In line with this, *TREM2* zf-hiMG in the *Irf8^st95^* background displayed a significantly smaller gastrosome compared to WT zf-hiMG in the same apoptotic context (FIG 3E, orange boxplots, rightmost vs. leftmost column; FIG 4B for a representative example). When transplanted into *slc37a2* mutant embryos, which retain endogenous microglia, *TREM2* zf-hiMG showed an even further reduction in gastrosome size, likely reflecting reduced access to apoptotic material due to competitive engulfment (FIG 3E, rightmost column, compare orange to pink boxplot). Despite their diminished phagocytic performance, *TREM2* zf-hiMG retained dynamic morphology and active scanning behavior, indicating that they remained viable and responsive *in vivo* (SUPP FIG 4D, Video 11). Together these findings demonstrate that the gastrosome in human microglia-like cells adapts to the rate of neuronal cell death and is sensitive to genetic perturbations in efferocytosis pathways. The observed reduction in gastrosome size in TRME2-deficient cells further underscores the role of this compartment in neuronal clearance and establishes the gastrosome as a conserved, functionally responsive structure central to microglial phagocytic competence.

**Figure 4.**
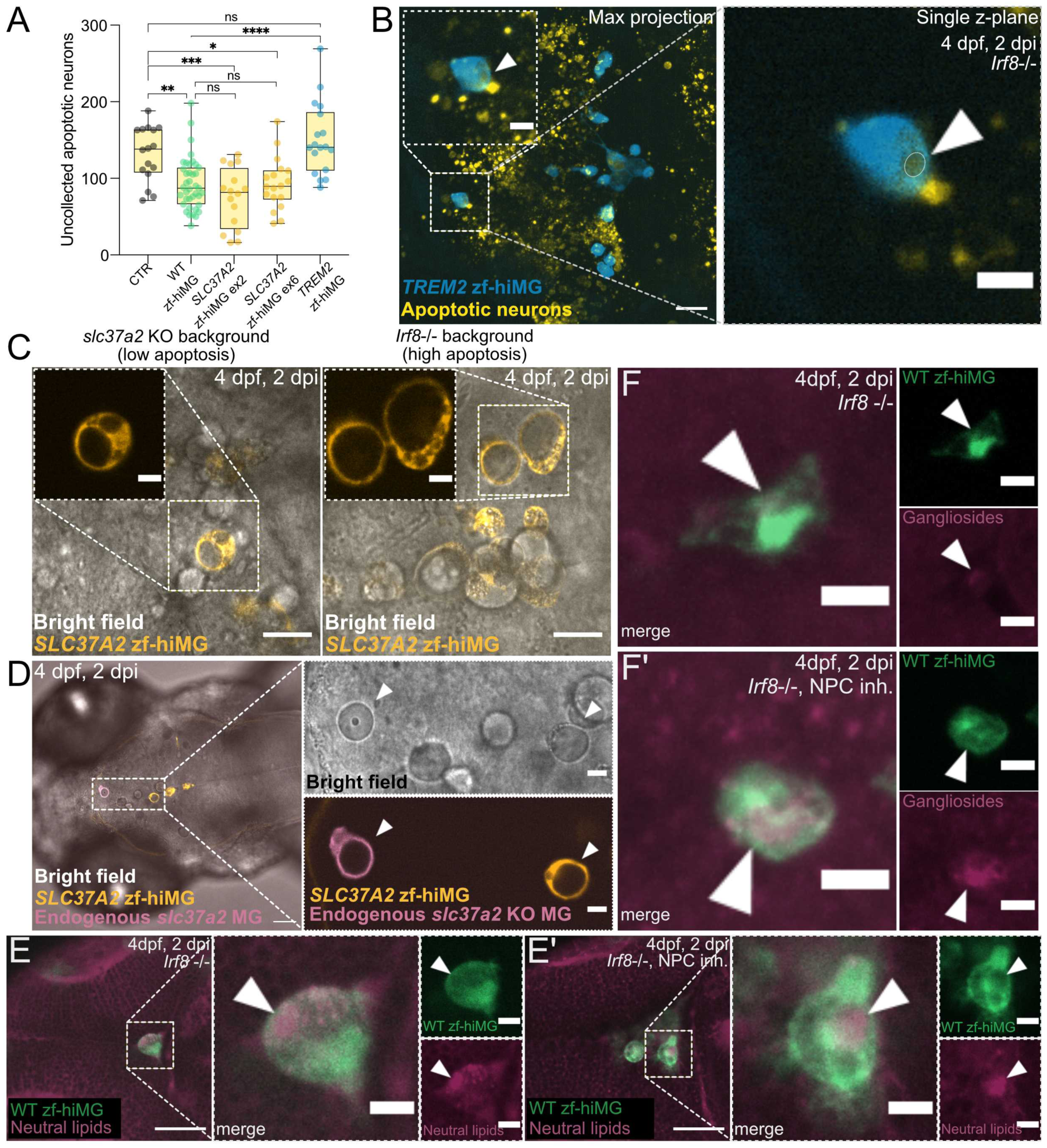
Genetic and functional perturbations alter gastrosome dynamics and lipid accumulation in human microglia-like cells *in vivo*. **(A)** Quantification of uncollected Acridine Orange (AO) positive apoptotic neurons in *Irf8^st95^* embryos without (CTR, grey dots) or with transplantation of WT (green dots), *SLC37A2* mutant (orange dots, exon 2 and exon 6 clones) or *TREM2* mutant (blue dots) zf-hiMG ; 3 dpf, 1dpi; N = 6 independent experiments, n (embryos) = 16 (CTR), 39 (WT zf-hiMG), 16 (*SLC37A2* ex2 zf-hiMG), 18 (*SLC37A2* ex6 zf-hiMG), 18 (*TREM2* zf-hiMG). **(B)** Representative image of GFP-labelled *TREM2* zf-hiMG in *Irf8^st95^* embryos with neuronal apoptotic marker (Tg(nbt:dLexPR-LexOP:secA5-BFP), arrowhead pointing at the largest vesicle in the cell; dorsal view; scale bars 30 µm (overview) and 10 µm (zoomed crop). **(C)** Bright field and fluorescence representative images of *SLC37A2* zf-hiMG in *Irf8^st95^* and *slc37a2* KO backgrounds; dorsal view, 4dpf, 2 dpi; scale bars 30 µm (overview) and 10 µm (zoomed crop). **(D)** GFP-labelled *SLC37A2* zf-hiMG into *slc37a2* KO embryos with red-labelled endogenous microglia; dorsal view, 4dpf, 2 dpi; scale bars 30 µm (overview) and 10 µm (zoomed crop). **(E-E’)** Neutral lipid staining (HCS LipidTox) in xenotransplanted GFP-labelled WT zf-hiMG in *Irf8^st95^*; with **(E’)** or without **(E)** pharmacological inhibition of NPC1; dorsal view; 4 dpf, 2 dpi; scale bars 50 µm (overview) and 10 µm (zoomed crop). **(F)** Staining with fluorescently labelled cholera toxin B (CtxB) to visualize GM1 gangliosides in GFP labelled WT zf-hiMG transplanted into *Irf8^st95^* with **(F′)** or without **(F)** NPC1 inhibition; 4 dpf, 2 dpi; dorsal view; scale bars 10 µm.

### In vivo characterization of the gastrosome in human microglia-like cells reveals a conserved role in lipids handling

To further explore the functional relevance of the gastrosome in human microglia, we next focused on its regulation and size. Given that *SLC37A2* has been implicated as a critical regulator of the gastrosome^13,14^, we used CRISPR-Cas9 genome editing to generate *SLC37A2*-deficient human iPSCs by targeting either exon 2 or exon 6 (SUPP FIG 4C). Both resulting clones harbored indels confirmed by sequencing (SUPP FIG 4C’) and showed a reduction in transcript levels by RNA-seq (SUPP FIG 4B). Pluripotency was retained in hiPSCs (data not shown) and the mutant cells were successfully differentiated into iMGL (*SLC37A2* iMGL) expressing canonical microglial markers at levels comparable to WT iMGL (SUPP FIG 4B). We transplanted *SLC37A2-*deficient iMGL into *Irf8^st95^* embryos and labeled apoptotic neurons using AO. Quantification revealed that both xenotransplanted *SLC37A2-deficient* lines significantly reduced apoptotic burden compared to non-transplanted controls, reaching levels similar to those achieved by WT zf-hiMG (FIG 4A). However, despite this preserved engulfment capacity, *SLC37A2-deficient* zf-hiMG showed a markedly enlarged gastrosome 48hrs post transplantation compared to WT zf-hiMG in the same background (FIG 3E, compare orange boxplots in middle vs. left columns; FIG 4C for representative images) indicating a defect in downstream processing of engulfed material. To further investigate this phenotype, we transplanted green-labelled *SLC37A2-defient* iMGL into *slc37a2* mutant embryos, in which endogenous microglia were labelled with mCherry. Two days post-transplantation, human and zebrafish *SLC37A2*-deficient microglia exhibited highly similar morphologies, appearing nearly indistinguishable (FIG 4D). Interestingly, the gastrosome size in *SLC37A2-deficient* zf-hiMG was smaller in *slc37a2* mutant embryos than in Irf8^st95^ embryos, likely reflecting reduced availability of apoptotic substrates due to the presence of resident microglia (FIG 3E, middle column; compare pink vs. orange boxplots; FIG 4C).

To assess the biochemical content of the enlarged gastrosome, we stained WT and *SLC37A2-deficient* zf-hiMG in the *Irf8^st95^* mutant background with cholera toxin B (CtxB), a marker for GM1 ganglioside. This revealed an accumulation of GM1 gangliosides within the enlarged gastrosome of *SLC37A2-deficient* zf-hiMG, indicating lipid buildup (SUPP FIG 4E). Lipid accumulation is a well-described feature of microglia in neurodegenerative disease contexts and is thought to reflect impaired degradation or trafficking of lipid-rich neuronal debris^5,6,9–12,14^. To test whether gastrosomal expansion reflects a general response to disrupted lipid trafficking, we transplanted human iMGL into *Irf8^st95^* embryos and exposed these embryos to U18666A, a well-characterized inhibitor of NPC1 -a cholesterol transporter whose mutation causes Niemann-Pick disease type C^51^. NPC1 is essential for proper microglial function, and microglial somatic enlargement has been reported in this disease context^14,52^. Upon NPC1 inhibition, we observed a robust expansion of the gastrosome in zf-hiMG mirroring the response of endogenous microglia within the same embryo (SUPP FIG 4F-F’). Staining with HCS LipidTOX and CtxB confirmed the accumulation of neutral lipids and gangliosides within the enlarged gastrosome (FIG 4E-E’ for neutral lipids and FIG 4F-F’ for gangliosides).

Together, these results demonstrate that the gastrosome is a conserved and functionally responsive compartment in human microglia. Its expansion in response to genetic and pharmacological disruption of lipid processing highlights the gastrosome’s essential role in the degradation and clearance of lipid-rich neuronal debris – a function critical for sustaining microglial homeostasis in both healthy and disease-associated states.

## Discussion

Impaired phagocytosis and disrupted lipid processing in microglia are increasingly recognized as contributing factors in the pathogenesis of neurodegenerative diseases. To investigate how human microglia process neuron-derived material, iPSC-derived microglia (iMGL) offer a powerful tool, providing a human genetic background and avoiding species-specific differences in gene regulation and behavior. However, to accurately capture microglial function, these cells must be studied in environments that mimic native tissue architecture and cues. Rodent xenotransplantation models and brain organoids provide valuable complexity and physiological relevance, but they are also time-consuming, technically demanding, and limited in throughput. In this context, HuZIBRA offers a complementary approach. Transplantation of iMGL into the developing zebrafish brain is technically simple, scalable, and allows for semi-throughput experimentation. While questions related to long-term integration of human microglia may be better suited to organoid or rodent systems, HuZIBRA can help with capturing acute, cell-autonomous microglial responses to neuronal apoptotic and metabolic stress. The zebrafish brain provides a permissive, in vivo environment in which human microglia exhibit highly branched morphologies and significantly more dynamic behavior than in vitro. Although some behavioral features may be dependent on the specific zebrafish context, the model’s small size and optical transparency enable high-temporal-resolution imaging of human microglia in real time. This reduces the risk of underestimating cell speed or oversimplifying migratory trajectories—limitations commonly encountered in slower or lower-resolution imaging platforms. Within HuZIBRA, human microglia are observed making transient, repeated contacts with dying zebrafish neurons. Such interactions resemble behaviors reported for endogenous zebrafish microglia ^32,37^, and echo transient synaptic contacts observed in rodent models^53^ (for review on the topic see^54^). Together, these findings point to a conserved feature of microglial surveillance and efferocytosis across species, highlighting HuZIBRA as a powerful tool for investigating core aspects of human microglial behavior in vivo. While the HuZIBRA model offers clear advantages in terms of scalability, accessibility, and imaging resolution, it also has some limitations. In particular, observations are typically limited to the first three days post-transplantation, when the embryos remain optically transparent—a key factor that dictates the feasible imaging window. Since human microglia do not proliferate during this time window, the model is less suited for applications that require large amounts of material, such as bulk or single-cell omics. However, this limitation is likely to become less relevant with the continued advancement of spatial transcriptomics—a rapidly evolving field to which HuZIBRA is well positioned to contribute in the future.

A key outcome of this study is the demonstration of conserved microglial efferocytosis across species. When transplanted into the zebrafish brain, human iPSC-derived microglia effectively recognize and engulf dying zebrafish neurons, indicating that core molecular cues guiding this process are functionally preserved. This cross-species compatibility provides a unique opportunity to model human microglial responses to neuronal cell death in vivo, and to dissect how genetic or pharmacological perturbations influence their clearance behavior. These dynamic processes are often understudied, partly due to the technical difficulty in imaging highly dynamic processes deep in the brain. Yet, they are highly relevant to understand how microglia respond to high levels of neuronal apoptosis that are typical of development and neurodegenerative conditions. To date, omics approaches have identified key gene expression programs and regulatory networks, however, our understanding of how these programs translate into core cellular processes, such as debris clearance, lipid metabolism, and dynamic environmental responses, remains limited. In this study, we observed that human microglia process apoptotic neuronal material through a coordinated pathway that involves the lipid-rich gastrosome. This compartment expands in response to increased neuronal death and is significantly reduced in microglia lacking functional TREM2, a receptor linked to neurodegenerative disease. These findings establish a mechanistic connection between gene expression programs and key cellular functions, namely efferocytosis and lipid metabolism, and underscore the value of HuZIBRA as a platform for dissecting both intrinsic and environmental regulators of microglial behavior in real time.

## Authors contribution

**A.V.** designed the study together with F.P, performed experiments, conducted data analysis, and contributed to conceptualization, writing, and editing of the manuscript. **T.W.** generated CRISPR-edited hiPSC lines. **J.W.** developed scripts for EM segmentation and provided feedback on data analysis. **I.M.G.** performed the bulk RNA-seq analysis. **I.S.O.** generated EM data of iMGL within human brain organoids. **N.T.** performed HCR experiments. **M.P.** contributed to EM data acquisition. **A.A.P** generated the Tg(UAS:lyn-miRFP670) fishline. **D.G.** provided feedback throughout the project, and manuscript review. **S.T.S.** supervised EM experiments and contributed to manuscript feedback. **F.P.** designed the study together with A.V. and contributed to conceptualization, writing, and editing of the manuscript.

## Acknowledgements

We are grateful to Cornelia Henkel for the fish care. The authors acknowledge the support of the Center for Microscopy and Image Analysis (ZMB), University of Zurich for light and electron microscopy experiments and of the Cytometry Facility for the support with FACS experiments. We are thankful to the Functional Genomic Center Zurich (FGCZ) of University of Zurich and ETH Zurich for the support performing the Genomics experiments. We are thankful to the Allen Institute for Cell Science for the generation and characterization of the labelled hiPSC lines. This project was supported by Swiss National Science Foundation Grants 310030_212794.

## Material and Methods

### Animal Model

Zebrafish (Danio rerio) were raised, maintained, and bred according to standard procedures as described in “Zebrafish – A practical approach” (Nüsslein-Volhard, 2012). All experiments were performed on embryos younger than 5dpf, in accordance with the European Union Directive 2010/62/EU and local authorities (Kantonales Veterinäramt; Fishroom licence TVHa Nr. 178). Live embryos were kept at 28°C in E3 solution and staging was done according to^55^. In this study sex determination is not relevant since in zebrafish all individuals develop initially immature ovaries, and the process of gonadal differentiation and sex determination takes place around 25 days post-fertilization^56^. To avoid pigmentation 0.003% 1-phenyl-2-thiourea was added at 1dpf. The following lines were used in this study *irf8^st95^* ^31^, *slc37a2^NY007^* ^13^, Tg(mpeg1:GFP-caax)^13^, Tg(nbt:dLexPR-LexOP:secA5-BFP)^32^, TgBAC(fms:Gal4,UAS:nfsB-mCherry)^57^, the Tg(UAS:lyn-miRFP670) plasmid was obtained by cloning the lyn-miRFP670 fusion protein downstream of the UAS promoter using the Gateway cloning kit (Thermo Fisher Scientific). This construct, flanked by Tol2 sites, was injected into 1-cell-stage Tg(fms:Gal4) embryos to generate transgenic lines expressing membrane-labeled, far-red microglia.

### Maintenance of hESC and iPSC and generation of fluorescnet reporter lines

Labelled human induced pluripotent stem cells (iPSC) AICS-0036-006 (WTC-mEGFP-Safeharborlocus(AAVS1)-cl6(mono-allelic tag)) and AICS-0054-091 (WTC-mTagRFPT-CAAX-Safeharborlocus(AAVS1)-cl91 (mono-allelic tag)) were developed at the Allen Institute for Cell Science (allencell.org/cell-catalog) and available through Coriell. The lines were cultured in mTeSR1 or mTeSR Plus (STEM CELL Technologies) in a humified incubator at 5% CO2, 37°C by following the guidelines provided by STEMCELL Technologies. All cell lines were regularly tested for mycoplasma contamination.

In experiments using hiPSC-derived microglia in brain organoids the protocol for the use of human embryonic stem cell lines (H1 ESC, WiCell) as well as human induced pluripotent stem cell (iPSC) lines for this study was approved by Salk Institute’s IRB Committee (FWA 00005316) and the Embryonic Stem Cell Research and Oversight Committee. The Salk Institute is committed to protecting the rights and welfare of human research participants and ensures compliance with all applicable ethical and legal requirements. The iPSC lines used in this study have previously been described^58^ and were reprogrammed in the same facility (Salk Institute for Biological Studies, Laboratory of Genetics) and under the same conditions. Briefly, fibroblasts were transduced with retroviruses containing SOX2, OCT4, KLF4 and MYC to induce overexpression of these genes and were transferred to a co-culture system with murine embryonic fibroblasts. iPSC colonies were identified after around 2 weeks in this culture system, plated onto Matrigel-coated plates (BD Biosciences) and maintained in iPS-Brew media (Miltenyi). All cell lines were regularly tested for mycoplasma contamination. The following line was used: Neurotypical control iPSC line Cent 3-6 as previously described^58^. To generate fluorescent reporter lines, we infected iPSCs with LV-CAG::tdT^23^. Five days after infection, cells were isolated using FACS and re-plated on Matrigel-coated plates in iPS-Brew media (Miltenyi) supplemented with CloneR (Stem Cell Technologies). Upon recovery, cell lines were expanded and used for subsequent experiments.

### CRISPR-mediated editing of SLC37A2 and TREM2

gRNA targeting the second exon of *TREM2* (5’-CTCTCCCAGCTGGCGGCACC-3’)^44^, or the second (5’-CTATCAGTATCGTCAAGGTG-3’) or sixth (5’-AATGGACTCGTCCAGACCAC-3’) exons of *SLC37A2* were designed using the CRISPR/Cas9 target online predictor CHOPCHOP^59^ and ordered from IDT as crRNA. The RNP complex was obtained following the manufacturer guidelines (IDT). Human iPSC were cultured in StemFlex Medium (ThermoFisher) and the RNP complex was introduced by electroporation (Lonza). Cells were diluted and seeded as single cells to obtain monoclonal lines using the isoCell-isoHub system (iota biosciences, Alameda US) by following the manufacturer guidelines. Successful targeting was confirmed by PCR and sequencing.

### Differentiation of iPSC-microglia (iMGL) from iPSCs

Human iPSC were differentiated to hemotopoietic progenitors (iHPC) using the Hematopoietic Kit (STEMCELL Technologies) and then to mature microglia (iMGL) using the Microglia Differentiation and Microglia Maturation kits (STEMCELL Technologies) by following the manufacturer guidelines.

### Generation of forebrain organoids

Subject-derived iPSC lines were used to generate forebrain organoids as described previously with minor modifications^58,60^. Human iPSC colonies were detached before reaching confluency with collagenase Type IV (Gibco) and transferred to an Ultra-Low attachment 10-cm plate (Corning Costar) containing 10 ml hPSC medium consisting of DMEM:F12 (Invitrogen), 20% Knockout Serum Replacement (Gibco), 1× Non-essential Amino Acids (Invitrogen), 1× 2-mercaptoethanol (Gibco), 1× GlutaMAX (Invitrogen), 10 ng ml–1 FGF-2 (Peprotech) and ROCK inhibitor Y27632 (10 μM). Twenty-four hours later, the medium was replaced with induction medium containing hPSC media without FGF-2, 2 μM dorsomorphin (Tocris) and 2 μM A-083 (Tocris). At day 5 the media was replaced with neural induction medium consisting of DMEM:F12 (Invitrogen), 1× N2 Supplement (Invitrogen), 1× Non-essential Amino Acids (Invitrogen), 1× GlutaMAX (Invitrogen), 10 μg ml–1 Heparin (Tocris), 1× Penicillin/Streptomycin (Gibco), 10 μM CHIR99021 (Tocris) and 1 μM SB-431542 (Tocris). Seven days after induction, organoids were embedded in 20-μl Matrigel (CultrexTM, Bio-Techne) droplets and continued to grow for an additional week in 6 cm Ultra-Low attachment plates (Corning Costar). From day 14 onwards, organoids were cultured in differentiation medium consisting of DMEM:F12 (Invitrogen), 1× N2 and B27 Supplements (Invitrogen), 1× Non-essential Amino Acids (Invitrogen), 1× GlutaMAX (Invitrogen), 1× 2-Mercaptoethanol (Gibco), 1× Penicillin/Streptomycin (Gibco) and 2.5 μg ml–1 Insulin (Sigma), and transferred to an orbital shaker (65-75 rpm). At day 20, residual Matrigel was removed and media changes were performed every 2-3 days using the aforementioned differentiation medium.

### Colonization of cortical forebrain organoids through erythromyeloid progenitors (EMPs)

EMPs were generated as previously described with minor modifications^29^. Briefly, hESCs/iPSCs were dissociated using TrypLE (Invitrogen) and plated in a 6-cm tissue culture-treated plate (CytoOne USA Scientific) at a density of 400,000 cells per well with 10 μM ROCK inhibitor (Tocris). The next day, cells were changed to basal hematopoietic differentiation media supplemented with FGF2 (50 ng ml-1, Miltenyi), BMP4 (50 ng ml-1, Proteintech) Activin A (12.5 ng ml-1, Miltenyi) ROCK inhibitor (1 μM, Tocris) and LiCl (2mM, Sigma) and grown under hypoxic conditions (5% CO2, 5% O2). Hematopoietic differentiation media consisted of IMDM (50%, Thermo Fisher Scientific), DMEM/F12 (50%), ITSG-X, (2% v/v, Thermo Fisher Scientific), L-ascorbic acid 2-Phosphate magnesium (64 mg ml-1; Sigma), monothioglycerol (400 mM), PVA (10 mg ml-1; Sigma), Glutamax (1X, Thermo Fisher Scientific), chemically defined lipid concentrate (1X, Thermo Fisher Scientific), non-essential amino acids (NEAA, Thermo Fisher Scientific 1X), Penicillin/Streptomycin (P/S Thermo Fisher Scientific, 1% V/V). On day 2, media was changed and supplemented with FGF2 (50 ng ml-1, Miltenyi) and VEGF (50 ng ml-1, Miltenyi) and returned to hypoxic conditions. Following day 4, cells were placed into norm-oxic conditions (5% CO2, 21% O2) and kept in basal hematopoietic differentiation media supplemented with FGF2 (50 ng ml-1, Tocris), VEGF (50 ng ml-1, Miltenyi) TPO (50 ng ml-1, Miltenyi), SCF (10 ng ml-1,Miltenyi), IL-6 (50 ng ml-1, Miltenyi) and IL-3 (10 ng ml-1, Miltenyi) until colonies released hematopoietic stem cells (usually between day 14 and 16). EMPs were then isolated using Fluorescence-activated cell sorting (FACS) by gating on tdT- and CD43-FITC (Biolegend, 315204)-positive cells. Co-culture of isolated EMPs with cortical organoids was performed using up to 100,000 cells in the above-indicated organoid differentiation media supplemented with 25 ng ml-1 M-CSF (Miltenyi), 100 ng ml-1 IL34 (Miltenyi) and 50 ng ml-1 TGFβ1 (Miltenyi).

### Flow cytometry

On day 12 of iHPC differentiation, hematopoietic markers were tested via antibody staining and FACS analysis. 5*10^4^ – 2*10^5^ cells/sample were incubated with antibodies diluted in FACS Buffer (D-PBS without Mg^++^ and Ca^++^ with 2% FBS) for 30min at 4°C. The cells were washed twice with FACS buffer and resuspended in 300ul sorting buffer (FACS buffer with 1mM EDTA and 25mM HEPES) for analysis. On day 24 of iMGL differentiation, microglial markers were tested via antibody staining and FACS analysis. 5*10^4^ – 2*10^5^ cells/sample were resuspended in 100 µl of FACS buffer and blocked with 100ul of 10µg/ml CD32 (20min at 4°C). The samples were then incubated 45 min at 4°C with the primary antibodies (see list below), and then 20 min at room temperature with secondary antibodies. The cells were washed twice with FACS buffer and then resuspended in 300ul sorting buffer for analysis. The samples were analyzed at a BD LSR II Fortessa Analyzer and processed in FlowJo.

### Phagocytosis assay *in vitro*

Mature iMGL seeded at a density of 1*10^5^ cells/cm^2^ on Matrigel-coated cell culture vessels were exposed to fluorescent Amyloid-β (Aβ) 1-42 (0.55ug/cm^2^), ovalbumin (4.21 ug/cm^2^) or 1µm large latex beads (diluted in a 1:5 ratio in not heat-inactivated FBS). 12-24 h after the feeding the cells were analyzed either by FACS or light microscopy.

### Staining procedures

#### HCR RNA-FISH *in vitro* and *in vivo*

The mRNA of selected targets was stained by HCR RNA-FISH (Molecular Instruments). HCR buffers, reagents, hairpins and probes were obtained from the manufacturer. The target mRNA sequences for the human-specific HCR probes were based on the MANE select transcript variants available on NCBI (Table 2). HCR probes were designed to reduce cross-reactivity between zebrafish and human genes (Table 5). For *in vitro* HCR experiments, mature iMGL were fixed by first adding 8% PFA directly to the culture medium to achieve a final concentration of 4% PFA (20 min at room temperature), followed by replacement with fresh 4% PFA in PBS which was incubated for 10 min at room temperature. HCR was then performed by following the manufacturer guidelines for mammalian cells on a chambered slide. For *in vivo* HCR experiments, 4 dpf zebrafish embryos were fixed overnight in PFA 4% at 4°C and dehydrated and permeabilized as by manufacturer guidelines with a proteinase K incubation of 30 minutes (protocol for whole-mount zebrafish embryos and larvae).

**Table 1.**
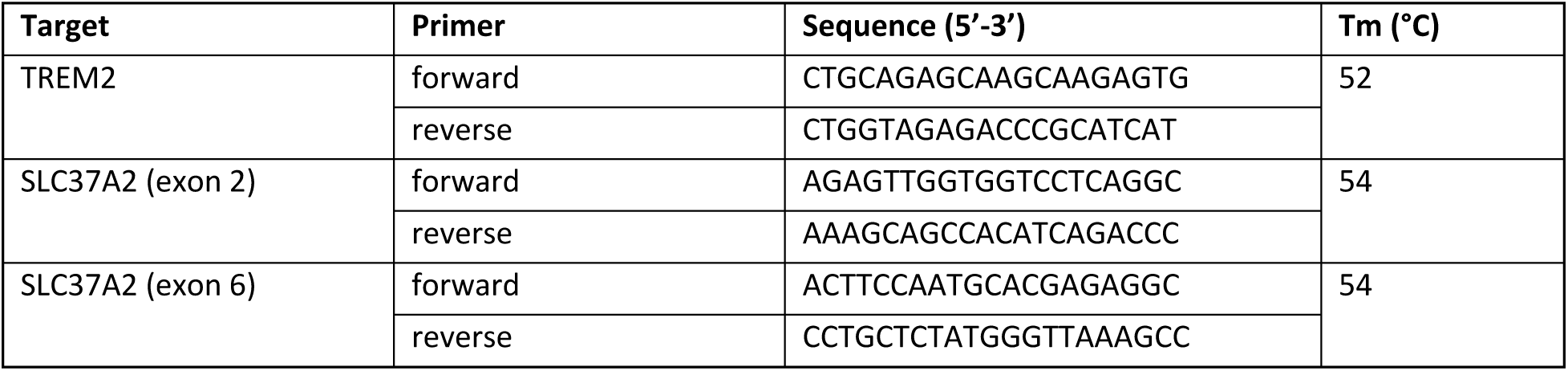
. List of primers for CRISPR sequencing.

**Table 2.** List of genes of interest, amplifiers and hairpins for HCR experiments.

| Gene of interest | NCBI accession number | HCR Amplifier | Hairpins |
| --- | --- | --- | --- |
| P2RY12 | NM_022788.5 | B2 | red ($\lambda$=594nm),<br>far red ($\lambda$=647nm) |
| AIF1 | NM_001623.5 | B1 | red ($\lambda$=594nm),<br>far red ($\lambda$=647nm) |
| TREM2 | NM_018965.4 | B1 | red ($\lambda$=594nm),<br>far red ($\lambda$=647nm) |
| TMEM119 | NM_181724.3 | B2 | red ($\lambda$=594nm),<br>far red ($\lambda$=647nm) |

#### Immunohistochemistry in cell culture

Mature iMGL were fixed by first adding 8% PFA directly to the culture medium to achieve a final concentration of 4% PFA (10-20 min at room temperature), followed by replacement with fresh 4% PFA in PBS which was incubated for 20 minutes at room temperature or overnight at 4°C. The cells were then rinsed in PBS and permeabilized for 7min at room temperature (0.2% Triton-X-100/PBS). After blocking for 1h at room temperature (3%BSA/0.2%Triton-X-100/PBS) the cells were incubated with the primary antibody diluted in blocking solution overnight at 4°C. The cells were rinsed 3 times with PBS1x and incubated 1h at room temperature with the secondary antibody and DAPI in blocking solution.

#### AnnexinV staining

A red labelled AnnexinV detection kit (ThermoFisher) was used to detect apoptosis *in vitro* by following the manufacturer guidelines. The samples were then analyzed by FACS. To analyse the staining by light microscopy, 3 µl of AnnexinV conjugate were added directly to a well of mature iMGL which were immediately imaged by spinning disk microscopy.

#### Acridine Orange

To visualize apoptotic cells *in vivo*, 3 dpf control and transplanted embryos were stained with acridine orange (AO, Merck, Sigma-Aldrich). The larvae were incubated in the dark for 1 h at 28 °C in 10 μg/ml AO solution in E3 and washed extensively before imaging.

#### GM1 ganglioside staining in zebrafish

To visualize GM1 gangliosides, a staining with Cholera Toxin subunit B (CtxB recombinant) conjugated with Alexa Fluor 549 or Alexa Fluor 647 (ThermoFisher Scientific) was performed on transplanted 4 dpf zebrafish embryos as previously reported (Zareba et al, 2024).

#### LipidTOX

To visualize neutral lipids, HCS LipidTOX Deep Red (ThermoFisher Scientific) was added 1:1000 to live 4dpf embryos for 1 hour at 28 degrees. The embryos were rinsed and mounted for live imaging.

### RNA sequencing

#### Library preparation

Total RNA was extracted from mature iMGL (1*106 cells) using either the RNeasy Micro kit (Qiagen) or the Direct-zol RNA miniprep (Zymo Research), according to the manufacturers’ protocols. The quality of the isolated RNA was determined with a Qubit® (1.0) Fluorometer (Life Technologies, California, USA) and a Fragment Analyzer (Agilent, Santa Clara, California, USA). Only those samples with a 260 nm/280 nm ratio between 1.8–2.1 and a 28S/18S ratio within 1.5–2 were further processed. The TruSeq Stranded mRNA (Illumina, Inc, California, USA) was used in the succeeding steps. Briefly, total RNA samples (100-1000 ng) were polyA enriched and then reverse-transcribed into double-stranded cDNA. The cDNA samples was fragmented, end-repaired and adenylated before ligation of TruSeq adapters containing unique dual indices (UDI) for multiplexing. Fragments containing TruSeq adapters on both ends were selectively enriched with PCR. The quality and quantity of the enriched libraries were validated using Qubit® (1.0) Fluorometer and the Fragment Analyzer (Agilent, Santa Clara, California, USA). The product is a smear with an average fragment size of approximately 260 bp. The libraries were normalized to 10nM in Tris-Cl 10 mM, pH8.5 with 0.1% Tween 20.

#### Cluster Generation and Sequencing

The Novaseq 6000 (Illumina, Inc, California, USA) was used for cluster generation and sequencing according to standard protocol. Sequencing was single end 100 bp.

#### Xenotransplantations of mature iMGL into Zebrafish embryos

2 days post fertilization (dpf) zebrafish embryos were anaesthetized and embedded ventrally (head up) as single embryos in small droplets of 1% low-melting agarose. The excess of agar was removed using a scalpel. Mature iMGL were harvested and filtered (mesh size 30-50 µm) after resuspension in HBSS with 1% Phenol Red to a final concentration of ∼2.4*10^6^ cells/ml. The cell suspension was loaded into a 30 µm beveled needle and injected directly into the optic tectum of the mounted embryos using an oil injector (Eppendorf). 10-20 minutes after injection, the embryos were unmounted and used for further experiments.

#### Light Microscopy

Confocal, spinning disk or light sheet microscopy were used to image the *in vitro* and *in vivo* samples, as specified below.

#### Light microscopy of cells

Confocal analysis of live or fixed cells was performed with an Andor Dragonfly 200 Sona spinning-disc microscope with a Nikon 20x/NA 0.95 water or 40x/NA 1.25 silicon oil objectives and 40 μm spinning disc or a Leica SP8 with a 20x/NA0.75 or 40x/NA1.1 water objectives to capture stacks with a z-step of 0.5-1 μm. All images were analyzed in Fiji^61^ or Imaris (Oxford Instruments).

#### Light microscopy of zebrafish embryos

Embryos were anaesthetized during mounting procedures and experiments using 0.01% tricaine (Merck) and pre-screened based on the expression of the desired fluorophore (when appropriate), using a Nikon ZMZ18 fluorescent stereoscope. Embryos were embedded in 1–1.5% low-melting (LM) agarose (PeqGOLD Low Melt Agarose, PeqLab Biotechnologie GmbH), dissolved in E3 medium with 0.01% tricaine and mounted as appropriate for the microscope used. For confocal microscopy, an Andor Dragonfly 200 Sona spinning-disc microscope with a Nikon 20x/NA 0.95 water or 40x/NA 1.25 silicon oil objectives and 40 μm spinning disc or a Leica SP8 with a 20x/NA0.75 or 40x/NA1.1 water objectives to capture stacks with a z-step of 0.5-1 μm. The same conditions were used for fixed zebrafish embryos. For light sheet microscopy we used either a Leica Viventis LS2 Live microscope with two Nikon 10x/0.2 NA objectives or a Bruker Luxendo TruLive3D Imager with two Nikon CFI Plan Fluor 10x/0.3 NA illumination objectives and a Nikon CFI APO LWD 25x/1.1 NA detection objective, coupled with a tube lens (f=400mm) to acquire images at 50x magnification. For time lapse images, the stacks were taken at an interval of 30 sec, 1 min, 3 min or 5 min. All images were analyzed in Fiji (Schindelin, J. et al., 2012) or Imaris.

### Chemical perturbations

#### U18666a

3dpf zebrafish embryos were incubated in 100 μM U18666a (Merck, Sigma-Aldrich) solution overnight at RT. Embryos were mounted and imaged.

#### Metronidazole

To induce apoptosis of endogenous zebrafish microglia, *slc37a2^NY007^* TgBAC(fms:Gal4,UAS:nfsB-mCherry) 3 dpf embryos were incubated with 10 mM Metronidazole (MTZ, Merck, Sigma-Aldrich) solution in E3 medium with 0.2% (v/v) DMSO for 24 h. Embryos were washed extensively with E3 before mounting and imaging.

#### Laser mediated injury

Injuries in the brain were performed at the Olympus IXplore SpinSR10 spinning disk confocal microscope equipped with the photomanipulation unit (Rapp OptoElectronic) using a 355 nm ablation laser (pulsed, UGA-42 Caliburn). For imaging, a 60x/1.3 NA objective was used, and for injury a circular pattern with 3% laser intensity was used.

### Electron Microscopy (EM)

#### TEM on iMGL

Transmission Electron Microscopy on iMGL (control or fed with latex beads) was performed at the Center for Microscopy and Image Analysis (ZMB) at the University of Zurich. Cells were grown as by differentiation protocol in 6 well plates and fixed with 2.5 % glutaraldehyde in 0.1 M sodium cacodylate buffer (pH 7.35, pre-warmed to 37°C), centrifugated at 3000 rpm for 6 min and washed in cacodylate buffer followed by 1% OsO4 for 1 hour in 0.1 M cacodylate buffer at 0°C, and 1% aqueous uranyl acetate for 1 hour at 4°C. Afterwards, cells were then first embedded in 2% agar, dehydrated in an ethanol series, followed by propylene oxide and embedded in Epon/Araldite (Sigma-Aldrich). Ultrathin (70 nm) sections were post-stained with lead citrate and examined with a Talos 120 transmission electron microscope at an acceleration voltage of 120 KV using a bottom mounted Ceta camera and the MAPS software for automatic image acquisition (Thermo Fisher Scientific, Eindhoven, The Netherlands).

#### Array Tomography on mature iMGL

Serial ultrathin sections (100nm) were collected on 10mm x 20mm silicon wafers using an ultramicrotome (Artos 3D, Leica Microsystems, Vienna, Austria)^62^ and contrasted with lead citrate for 7 min.

Sections were imaged in an Apreo 2 VS scanning electron microscope using the MAPS software package for automatic serial section recognition and image acquisition (Thermo Fisher, Eindhoven, The Netherlands). The array tomography workflow includes serial sections recognition, image region definition, autofunctions, and image acquisition^63^. Region of interest was imaged on every section using the OptiPlan mode of the system and the T1 detector with following parameters: Pixel size of 4 nm, pixel dwell time of 1 us, electron high tension of 1.8 keV, beam current of 0.1 nA. Serial section Tiff images were aligned using the plugin TrakEM2^64^ in Fiji^61^.

### Electron microscopy on human iPSC-microglia in human brain organoids

#### Fixed sample processing

Microglia-containing forebrain organoids were fixed at 6/7 weeks post integration using 2% PFA / 2% Glutaraldehyde in 0.1M Phosphate Buffer (pH= 6.8) for 2 hours at room temperature, followed by two washes in 0.1M Phosphate Buffer (pH= 6.8). The fixed organoid samples were rinsed in 0.1M cacodylate buffer and the localization of fluorescently labeled cells was determined by widefield fluorescence microscopy for tdTomato. The sample was microdissected for the fluorescently labeled region and embedded in 4% low melting point agarose made with 1X PBS. A Leica VT1000 vibratome was used to collect 100µm sections of the sample. Sections were stored in cryoprotectant (4:3:3 - PBS:ethylene glycol:glycerol) at −20°C until further processing. The sample was imaged on a Zeiss 880 microscope with a 20X air objective (NA 0.8) using the Airyscan detector. A comprehensive map of all tdTomato+ cells was generated using tile scanning and z-stack modules (reporting pixel size and volume size of whole ROI and sub-ROIs).

#### vEM sample preparation

Materials were sourced from Electron Microscopy Sciences (Hatfield, PA) unless otherwise stated. Samples were rinsed repeatedly with ice-cold buffer (0.1M sodium cacodylate, 3mM calcium chloride) before further fixation with reduced osmium (1% osmium tetroxide, 1.5% potassium ferrocyanide, 0.1M sodium cacodylate, 3mM calcium chloride). Samples were rinsed repeatedly with ice-cold water and stained with 1% aqueous uranyl acetate for one hour. Samples were rinsed thoroughly with ice cold water and serially dehydrated in ascending concentrations of ice-cold ethanol. Samples were finally rinsed in three changes of anhydrous ethanol at room temperature before infiltration with a 1:1 mixture of anhydrous ethanol and epoxy resin (Eponate 12, hard formulation; Ted Pella) for 4 hours on a rotating mixer, followed by infiltration in pure resin overnight on a rotating mixer. The samples were embedded in two steps: first, the infiltrated sample was polymerized overnight at 70°C on polypropylene bottlecap in a thin layer of resin with a gel capsule pressed into the resin around the sample. The following day, the gel capsule was backfilled with more resin and left to polymerize for another 48 hours.

Serial ultrathin sections (100nm) were collected from the blockface. Briefly, the blockface was trimmed using a 90° diamond trimming knife (Diatome) around the entire tissue section to a frustum of approximately 80µm in height. A silicon chip (35×7mm; University Wafer, Boston, MA) was hydrophilized in a plasma cleaner (Harrick), rinsed in pure water, and partially immersed in a Diatome Histo knife, with one end sticking out of the water at the back of the boat. Four drops of pure ethanol were added to the water in the boat to attenuate surface tension, and an ionizing gun (Leica EM Crion) was activated and oriented towards the cutting edge of the knife mounted on the ultramicrotome. Ribbons of approximately 100-150 serial sections of a nominal 100nm thickness were collected onto a series of chips, for a total of approximately 650 sections or 65µm of tissue volume. When ribbons of sufficient quality and length were generated, they were released from the knife edge using a single-eyelash brush and carefully positioned over the chip. The water level was then slowly lowered, and sections were allowed to dry down on the silicon substrate over a few minutes. Chips were further dried on a hot plate set to 60°C for approximately 5 minutes.

#### vEM imaging

Samples were loaded into a Zeiss Sigma VP scanning EM microscope (SEM) and imaged the array tomography software module of Atlas5 (FIBICS). Images were collected using a backscattered electron detector (Gatan) at a working distance of approximately 6mm, with accelerating voltage set to 3kV, a 30µm aperture, and the beam in high current mode. Low magnification (50-100nm/px) overviews of each section were collected and aligned using TrakEM2^64^ in Fiji^61^. These stacks were used for correlation with light microscopy, and subsequent selection of regions of interest for high resolution vEM. Correlative light-electron microscopy (CLEM) was achieved using BigWarp^65^ in Fiji^61^ between the tdTomato fluorescence and the microglia in the vEM. In sparse vEM regions, the only cells present were microglia, which had stereotypical ultrastructural features (eg., electron dense ER, large clear endosomes, and complexes formed from lipid droplets). The presence of these distinguishing features supported the positive identification of tdTomato+ microglia in the vEM data from more dense cellular regions.

#### EM Segmentation and visualization

Regions of interest including positively identified tdTomato+ microglia were imaged at high resolution (8nm/px) through continuous serial sections and aligned as described above. For segmentation of human cells in *in vitro* monoculture, first EM image stacks were registered using the StackReg FIJI plugin^66^. For segmentation we used the scikit-image toolkit^67^. For distinguishing between cytosol, background and gastrosome, we thresholded based on the mean intensity of SLIC super-pixels^68^ (parameters: n_seg=20000, compactness_slic=0.2, sigma_slic=4, min_size_factor_slic=0.5, max_size_factor_slic=3). For segmenting the nucleus, we used a seeded watershed, the seeds were manually defined in napari^69^ and interpolated between layers. Rough masks were created in napari to exclude neighouring cells. The resulting segmentation was checked in napari and manually improved it where appropriate. For segmentations of human iPSC-derived microglia in brain organoids, because of the lower contrast of this data set, the slices of the EM stack were segmented manually in napari. After segmentation, in both cases the resulting masks were imported in Imaris (RRID:SCR_007370) for visualisation purposes.

### Data Analysis

#### RNA sequencing

In-house RNA-seq raw reads were processed using ARMOR (PMC6643886). Reads were aligned and counted against the human genome GRCh38 assembly and Gencode release 43 annotation using salmon v1.4.0 and STAR 2.7.7a. Abud’s and iPSC data were downloaded from the Sequence Read Archive (SRA) accessions SRP092075 and SRP155574, respectively, using recount3 (PMID 34844637), with GRCh38 as the reference genome and Gencode’s annotation. The salmon-generated in-house count data were modelled with quasi-likelihood (QL) negative binomial generalized log-linear models, and differential expression analysis was performed using edgeR v3.36.0 on salmon outputs. Multi-dimensional scaling (MDS) plots were generated based on normalized count data using edgeR v3.36.0. Briefly, MDS plots were used to depict similarities between samples (including replicates and batches) in an unsupervised manner, showing the two leading fold-change dimensions which explain the largest proportion of variation in gene expression across samples. Batch correction was performed using ComBat from the R package sva v3.42.0, with the origin (Abud’s or in-house) used as the batch variable. Gene expression plots were generated to depict logCPMs as calculated by edgeR’s cpm(assay(x, ‘counts’), log = TRUE, prior.count = 2) or scaled and centered logCPMs, as reported in figure legends. Gene set enrichment analyses were performed using camera from limma v3.50.3 (PMC3458527) and mSigDB annotations on differentially expressed genes as reported by edgeR. Selected gene sets included: C2, curated gene sets from online pathway databases, publications in PubMed, and domain expert knowledge; C5, Gene Ontology; and C8, cell type signatures.

#### Transplants statistics

For each transplantation experiment, the total number of injected embryos was recorded. All embryos were imaged, and transplantation efficiency was calculated as the percentage of embryos showing zf-hiMG in the optic tectum (i.e., transplanted and correctly localized) over the total number injected. These embryos were then used to quantify the number of transplanted cells per embryo by manual counting in Fiji^61^. For survival analyses, embryos were repeatedly imaged from 2 days post-fertilization (dpf) to 5 dpf (corresponding to 0 to 3 days post-injection, dpi). Embryos were unmounted and remounted daily. Survival was expressed as the percentage of remaining cells over time, normalized to the initial number of cells at 0 dpi.

#### Morphology analysis

Morphological features of iMGL and zf-hiMG were manually analyzed on 2D maximum intensity projections using Fiji^61^. The area of the cell body was measured using the segmented line tool. To estimate the total area covered by a cell, the convex hull was manually drawn by connecting the distal tips of the branches to form the smallest possible polygon encompassing the entire cell. Additionally, the number of primary branches directly emerging from the cell body was manually counted.

#### Branch motility

Branch dynamics were analyzed on 2D maximum intensity projections from light-sheet microscopy data. When cells were migrating, the imaging frame was cropped to center the cell. Raw 3D volumes were converted to H5 format using Fiji BigDataProcessor^70^. Individual branches were tracked manually using the mTrackJ plugin^71^ in Fiji^61^, both *in vitro* and *in vivo*.

#### Cell body motility

Cell body motility was quantified in 2D using using the mTrackJ plugin^71^ in Fiji^61^. For the qualitative representation of zf-hiMG cell body motility, microglia were tracked semi-automatically using Imaris spot detection over time. Spot detection was automated and manually corrected to ensure accurate placement and tracking throughout the time-lapse.

#### Acridine Orange (AO) quantification

Quantification of uncollected apoptotic neurons was performed using acridine orange (AO) staining (described above) and Imaris spot detection. 3D volumes were analyzed to detect AO⁺ spots within the optic tectum, using a spot diameter of 4 μm. Automated detection was manually corrected. In transplanted samples, the segmentation channel was duplicated and overlaid with the zf-hiMG channel to exclude AO⁺ spots located within zf-hiMG. In experiments with green zf-hiMG, AO⁺ clusters recognizable as single, intense aggregates were excluded from quantification, as they were considered internalized by microglia. This exclusion procedure was applied when needed to both WT and mutant groups.

#### Fluorescence intensity

To confirm colocalization of apoptotic material within vesicular compartments (e.g., phagosomes, gastrosomes), fluorescence intensity profiles were extracted in Fiji^61^. A line was drawn across the vesicle of interest, and intensity values for both the zf-hiMG signal and the apoptotic neuron signal were recorded and overlayed.

#### Gastrosome diameter

The diameter of gastrosomes was quantified manually in Fiji^61^ on 3D image stacks. For each cell, the entire volume was scanned, and the largest vesicle was identified. Using the straight line tool, the longest possible axis within the vesicle was measured and recorded as the gastrosome diameter. Only the largest vesicle per cell was included in the analysis.

#### Statistical Analysis

Statistical analyses were performed using Prism 9 (GraphPad), Python, or R. Unless otherwise specified, conditions were compared using an unpaired, two-tailed, nonparametric Mann-Whitney U test with Bonferroni correction for multiple comparisons. The following p-value thresholds were used to report statistical significance:

**Table 2.** Summary of Statistical Significance Thresholds and Reporting Symbols.

| p-value | Wording | Summary |
| --- | --- | --- |
| < 0.0001 | Very highly significant | **** |
| 0.0001 – 0.001 | Highly significant | *** |
| 0.001 – 0.01 | Very significant | ** |
| 0.01 – 0.05 | Significant | * |
| ≥ 0.05 | Not significant | ns |

### Key resource tables

#### List of Materials

**Table 3.** Materials.

| Reagent type (species) or resource | Designation | Source or reference | Identifiers | Additional information |
| --- | --- | --- | --- | --- |
| Genetic reagent (*Danio rerio*) | *irf8<sup>st95</sup>* | Shiau, C. E. et al, 2015 | ZDB-FISH-150901-4256 |  |
| Genetic reagent (*Danio rerio*) | *slc37a2<sup>NY007</sup>* | Villani et al, 2019 | ZDB-FISH-191226-5 |  |
| Genetic reagent (*Danio rerio*) | Tg(mpeg1:GFP-caax) | Villani et al., 2019 | ZDB-TGCONSTRUCT-191211-1 |  |
| Genetic reagent (*Danio rerio*) | Tg(nbt:dLexPR-LexOP:secA5-BFP) | Mazaheri et al., 2014 | ZDB-TGCONSTRUCT-170110-1 |  |
| Genetic reagent (*Danio rerio*) | TgBAC(fms:Gal4,UA S:nfsB-mCherry) | Gray et al., 2011 | ZDB-ALT-110707-2 |  |
| Genetic reagent (*Danio rerio*) | Tg(UAS:lyn-miRFP670) | This paper | NA |  |
| Cell line | TC-mEGFP-Safeharborlocus(AA VS1)-cl6(mono-allelic tag) | Allen Institute for Cell Science | AICS-0036-006 |  |
| Cell line | WTC-mTagRFPT-CAAX-Safeharborlocus(AA VS1)-cl91 (mono-allelic tag) | Allen Institute for Cell Science | AICS-0054-091 |  |
| Commercial assay or kit | MultiSite Gateway Pro | ThermoFisher Scientific | 12537100 |  |
| Commercial assay or kit | STEMdiff Hematopoietic Kit | STEMCELL Technologies | 05310 |  |
| Commercial assay or kit | STEMdiff Microglia Differentiation Kit | STEMCELL Technologies | 100-0019 |  |
| Commercial assay or kit | STEMdiff Microglia Maturation Kit | STEMCELL Technologies | 100-0020 |  |
| Commercial assay or kit | Annexin V |  |  |  |
| Commercial assay or kit | RNeasy Micro kit | Qiagen | 74004 |  |
| Commercial assay or kit | Direct-zol RNA miniprep | Zymo Research | R2050 |  |
| Reagent | mTeSR1 | STEMCELL Technologies | 85850 |  |
| Reagent | MTeSR Plus | STEMCELL Technologies | 100-0276 |  |
| Reagent | ReLeSR | STEMCELL Technologies | 05872 |  |
| Reagent | Matrigel hESC-Qualified Matrix | Corning | 354277 |  |
| Reagent | Matrigel Growth Factor Reduced | Corning | 354230 |  |
|  | (GFR) Basement Membrane Matrix |  |  |  |
| Reagent | Y-27632 (Rock-Inhibitor) | STEMCELL Technologies | 05888 |  |
| Reagent | Accutase | STEMCELL Technologies | 07922 |  |
| Reagent | Alt-R® CRISPR-Cas9 crRNA | IDT |  |  |
| Reagent | Alt-R® CRISPR-Cas9 tracrRNA | IDT | IDT (1072533) |  |
| Reagent | Alt-R® S.p. HiFi Cas9 Nuclease V3 (stock 61uM) | IDT | IDT (1081060) |  |
| Reagent | Nuclease-free IDTE | IDT | IDT (11-01-01-01) |  |
| Reagent | StemFlex™ Medium | ThermoFisher Scientific | A3349401 |  |
| Reagent | Human Stem Cell Nucleofector™ Kit 1 | Lonza | VPH-5012 |  |
| Reagent | Alt-R® Cas9 Electroporation Enhancer | IDT | 1075915 |  |
| Reagent | CloneR | STEMCELL Technologies | 05888 |  |
| Reagent | TrypLE Express Enzyme (1X), no phenol red | ThermoFisher Scientific | 12604013 |  |
| Reagent | Acridine Orange | Merck (Sigma-Aldrich) | **A6014** |  |
| Reagent | CtxB 549 | ThermoFisher Scientific | C22842 |  |
| Reagent | CtxB 647 | ThermoFisher Scientific | C34778 |  |
| Reagent | HCS LipidTOX Deep Red | ThermoFisher Scientific | H34477 |  |
| Reagent | Tricaine | Merck | A5040 |  |
| Reagent | U18666a | Merck (Sigma-Aldrich) | U3633 |  |
| Antibody | Anti-human CD43 | STEMCELL Technologies | 60085AZ | 1:100 |
| Antibody | Anti-human CD45 | STEMCELL Technologies | 60018AZ.1 | 1:100 |
| Antibody | Anti-human CD34 | ThermoFisher Scientific | CD34-581-04 | 1:100 |
| Antibody | Anti-human CD45 | STEMCELL Technologies | 60018AZ.1 | 1:200 |
| Antibody | Anti-human CD11b | STEMCELL Technologies | 60040PE.1 | 1:100 |
| Antibody | Anti-human CD14 | STEMCELL Technologies | 60004AZ.1 | 1:200 |
| Antibody | Anti-human TREM2 | R&D Systems | AF1828 | 1:20 (FACS),<br>10ug/ml (IHC) |
| Antibody | Anti-human TMEM119 | Abcam | ab185333 | 1:500 (FACS),<br>1:1000 (IHC) |
| Antibody | Anti-human pU.1 | Cell Signaling | 2266S | 1:40 (FACS), 1:1000 (IHC) |
| Antibody | Anti-human P2Y12 | Merck | HPA014518 | 1:25 |
| Antibody | Anti-human IBA1 | Synaptic Systems | 234003 | 1:250 (FACS), 1:1000 (IHC) |
| Antibody | Anti-goat AF546 | ThermoFisher Scientific | A-11056 | 1:500 |
| Antibody | Anti-rabbit AF568 | Molecular Probes | A11011 | 1:500 |
| Instrument | Nucleofector™ 2b | Lonza | AAB-1001 |  |
| Instrument | IsoCell-isoHub | IotaSciences |  |  |
| Instrument | BD LSR II Fortessa Analyzer |  |  |  |
| Instrument | CellTram 4r Oil | Eppendorf |  |  |
| Instrument | Andor Dragonfly 200 | Andor |  |  |
| Instrument | SP8 | Leica |  |  |
| Instrument | Vivantis LS2 Live | Leica |  |  |
| Instrument | TruLive3D Imager | Brucker, luxendo |  |  |
| Instrument | IXplore SpinSR10 | Olympus |  |  |
| Software | CHOPCHOP |  | Labun, K. et al, 2019 |  |
| Software | FlowJo |  |  |  |
| Software | Fiji |  |  |  |
| Software | Imaris | Oxford Instruments |  |  |

#### List of CRISPR sequences

**Table 4.**
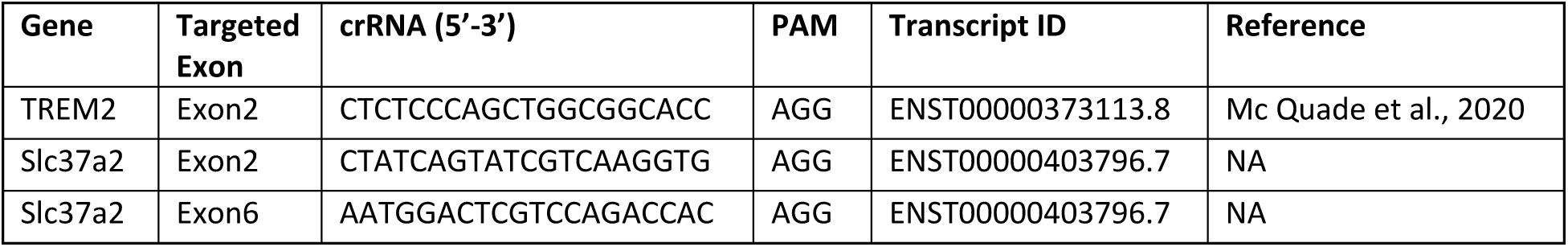
List of CRISPR sequences.

#### List of HCR probes sequences

**Table 5.**
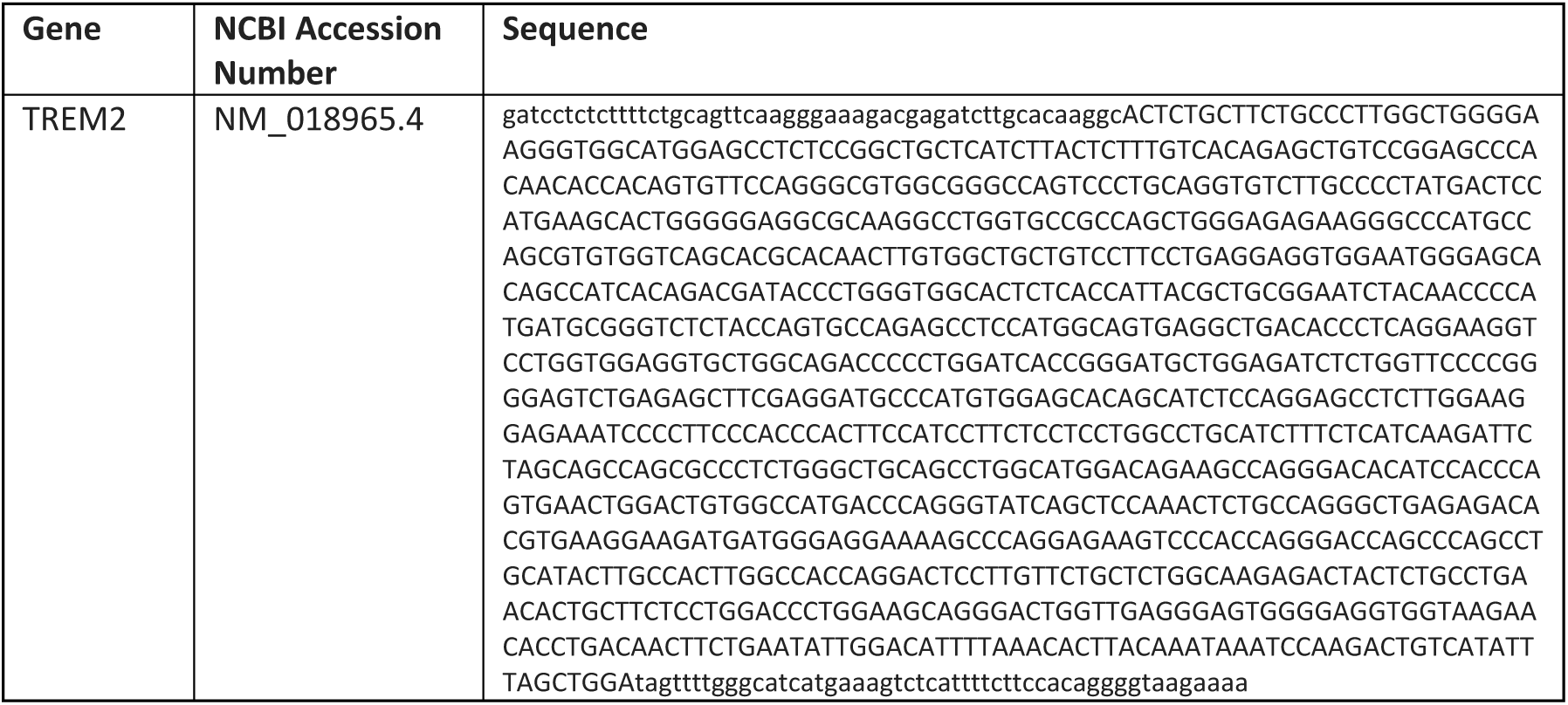
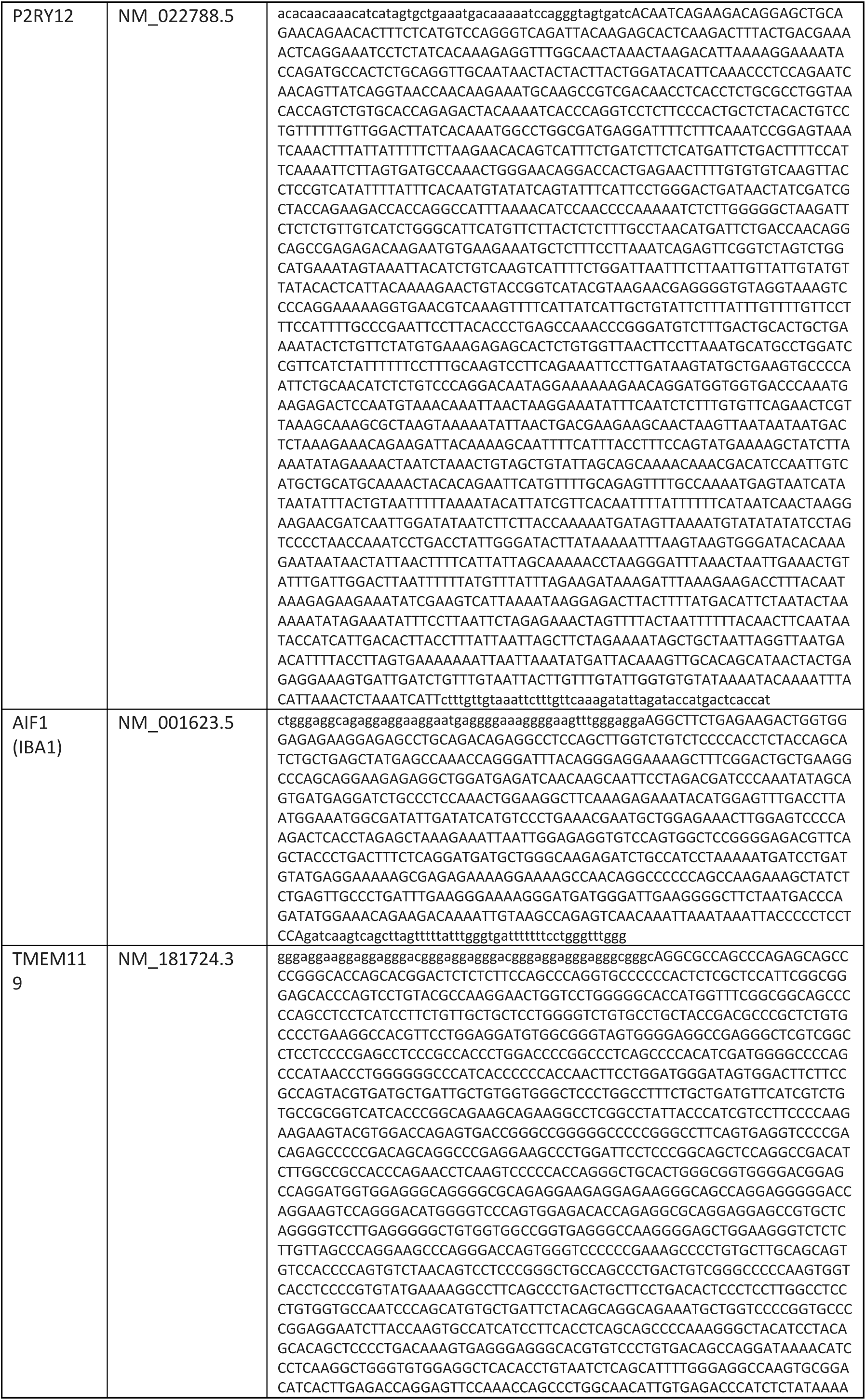
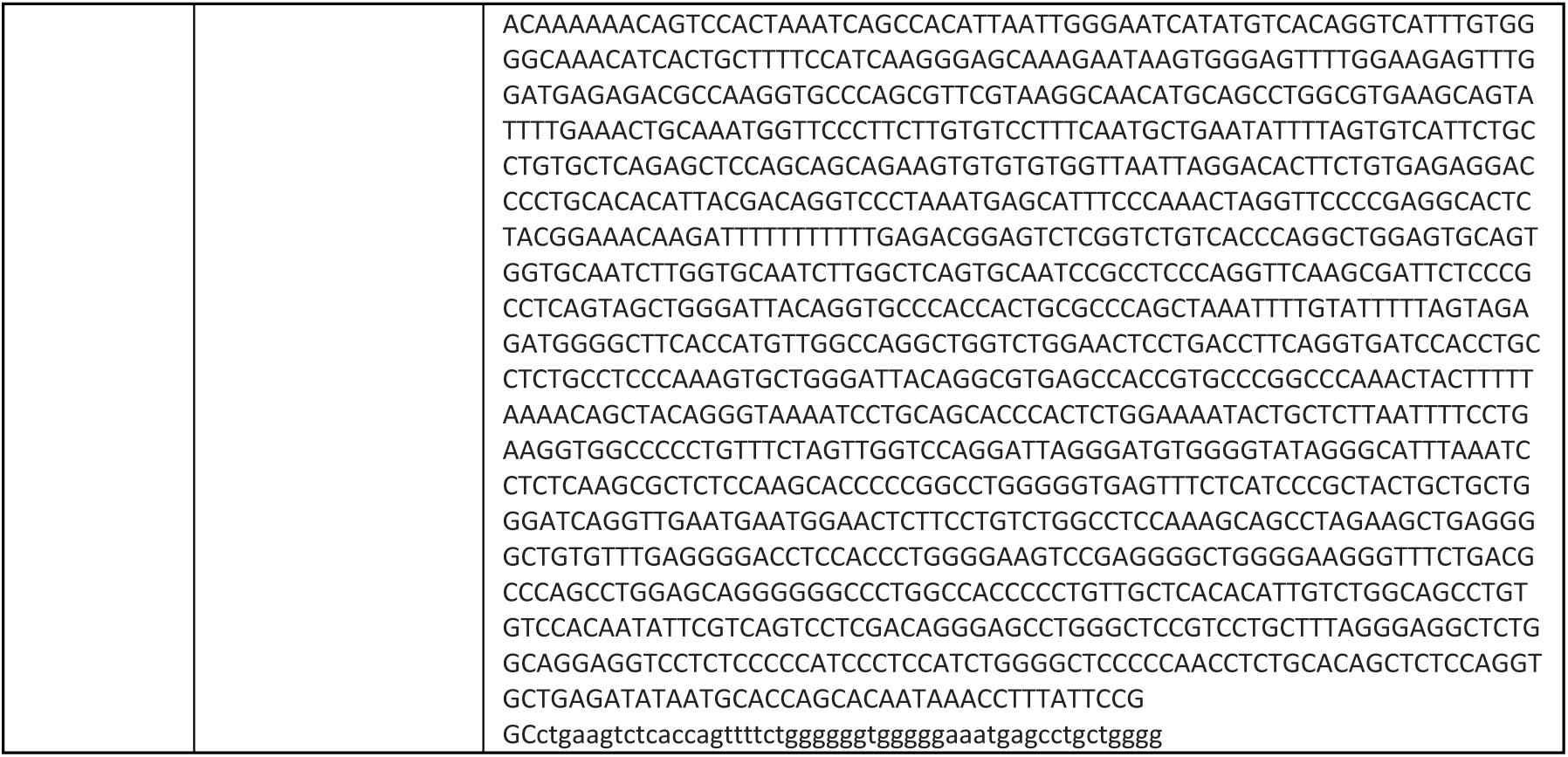
List of HCR sequences.

---

**Supplementary Figure 1.**
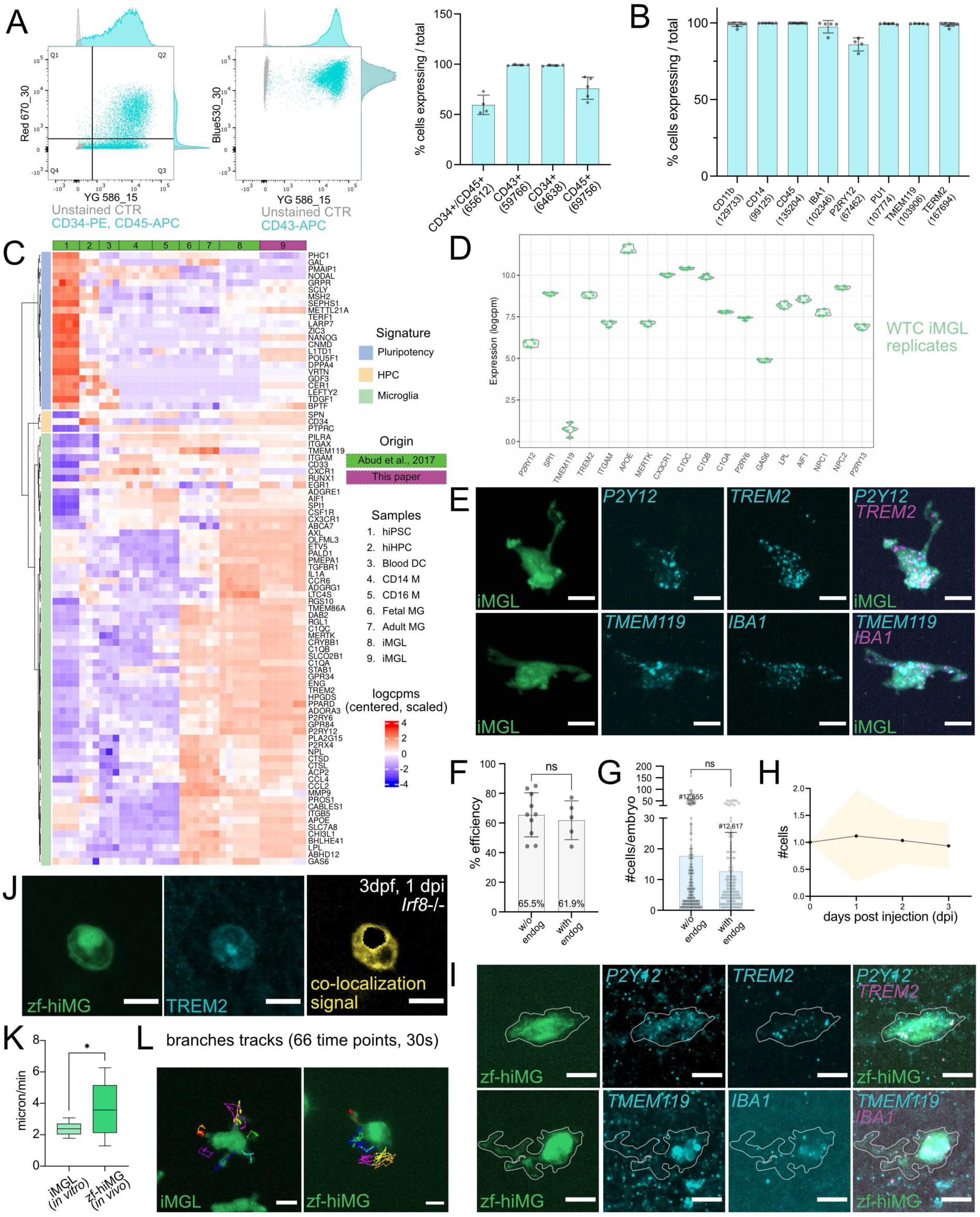
Characterization and validation of iMGL and their engraftment in zebrafish (zf-hiMG). **(A)** Flow cytometry analysis of iHPC generated from fluorescently labelled hiPSC, based on typical HPC marker expression; N=5 (experiments), number of cells in x axes. **(B)** Histogram showing FACS quantification of microglial protein markers in iMGL; N=9 (experiments), number of cells in x axes. **(C)** Heatmap of bulk RNA-seq data comparing WT iMGL generated in this study to published datasets of iMGL and other cell types (Abud et al. 2019). **(D)** Violin plots showing normalized expression levels of canonical microglial genes in WT iMGL. **(E)** *In vitro* Hybridization Chain Reaction (HCR) of key microglial transcripts in WT iMGL; scale bars 10 µm. **(F–G)** Quantification of xenotransplantation efficiency. **(F)** Percentage of *Irf8^st95^* embryos containing fluorescently labelled iMGL in the Optic Tectum (OT); N=10, n=530 (w/o endogenous); N=5, n=190 (with endogenous). **(G)** Number of zf-hiMG per embryo; N=8, n=171 (w/o endogenous); N=5, n=107 (with endogenous). **(H)** Quantification of zf-hiMG every 24hrs, from 2 dpf, 0 dpi to 5 dpf, 3 dpi; N=2, n=56. **(I)** *In vivo* HCR on zf-hiMG of human-specific microglial transcripts TMEM119, P2Y12, IBA1, and TREM2; scale bars 10 µm. **(J)** *In vivo* immunofluorescence staining of human TREM2 in zf-hiMG; scale bar 10 µm. **(K–L)** Analysis of zf-hiMG motility (*in vivo*) compared to iMGL (*in vitro*). **(K)** Quantification of branch extension/retraction speed *in vitro* versus *in vivo*; N=3, n=10 (*in vitro*), n=7 (*in vivo*). **(L)** Qualitative tracking of branch motility over time in representative iMGL (left) and zf-hiMG (right); scale bars 10 µm. N=experiments, n=embryos (F-H)/cells (K).

**Supplementary Figure 2.**
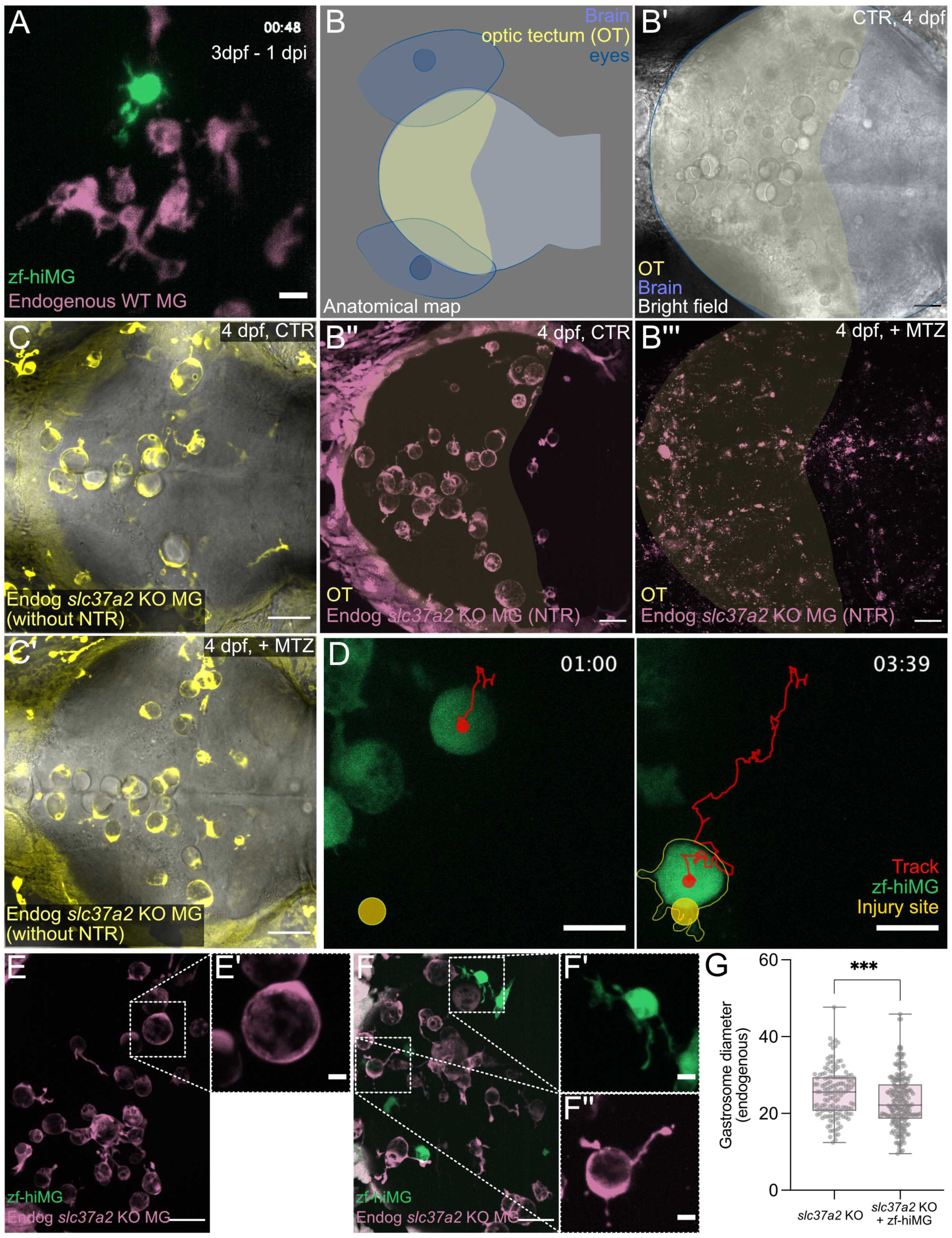
Coexistence and functional complementation of human and zebrafish microglia in vivo. **(A)** GFP labelled WT zf-hiMG transplanted into WT zebrafish embryos with red-labelled endogenous microglia (Tg(fms:Gal4;UAS:nfsB-mCherry)), OT dorsal view, 3 dpf, 1 dpi; 3 minutes time resolution; scale bar 10 µm. **(B)** Dorsal view anatomical map of a zebrafish embryo showing segmented regions of the eyes, brain, and optic tectum (OT); **(B’)** Bright field image of a control *slc37a2* KO embryo at 4 dpf; **(B’’)** Image of control *slc37a2* KO endogenous microglia (Tg(fms:Gal4;UAS:nfsB-mCherry)) at 4 dpf; **(B’’’)** Image of MTZ-treated *slc37a2* KO endogenous microglia (Tg(fms:Gal4;UAS:nfsB-mCherry)) at 4 dpf; scale bars 50 µm. **(C–C’)** Fluorescence and bright field images of *slc37a2* KO embryos at 4 dpf lacking NTR expression (Tg(mpeg1:eGFP-caax)) without **(C)** or with **(C’)** MTZ treatment; scale bar 50 µm. **(D)** Injury response of GFP-labelled WT zf-hiMG in *Irf8^st95^* 4 dpf, 2 dpi zebrafish embryo; 5 minutes time resolution; scale bar 10 µm. **(E–F)** Dorsal views of *slc37a2* KO 4 dpf embryos (Tg(fms:Gal4;UAS:nfsB-mCherry)); scale bars 50 µm (overviews) and 10 µm (cropped zooms): **(E)** without zf-hiMG transplant; **(F)** with zf-hiMG transplant (2 dpi). **(G)** Quantification of E-F: gastrosome diameter of endogenous *slc37a2* KO microglia ± zf-hiMG transplantation; N=7, n=138 (*slc37a2* KO); N=9, n=202 (*slc37a2* KO+zf-hiMG); N=embryos, n=cells.

**Supplementary Figure 3.**
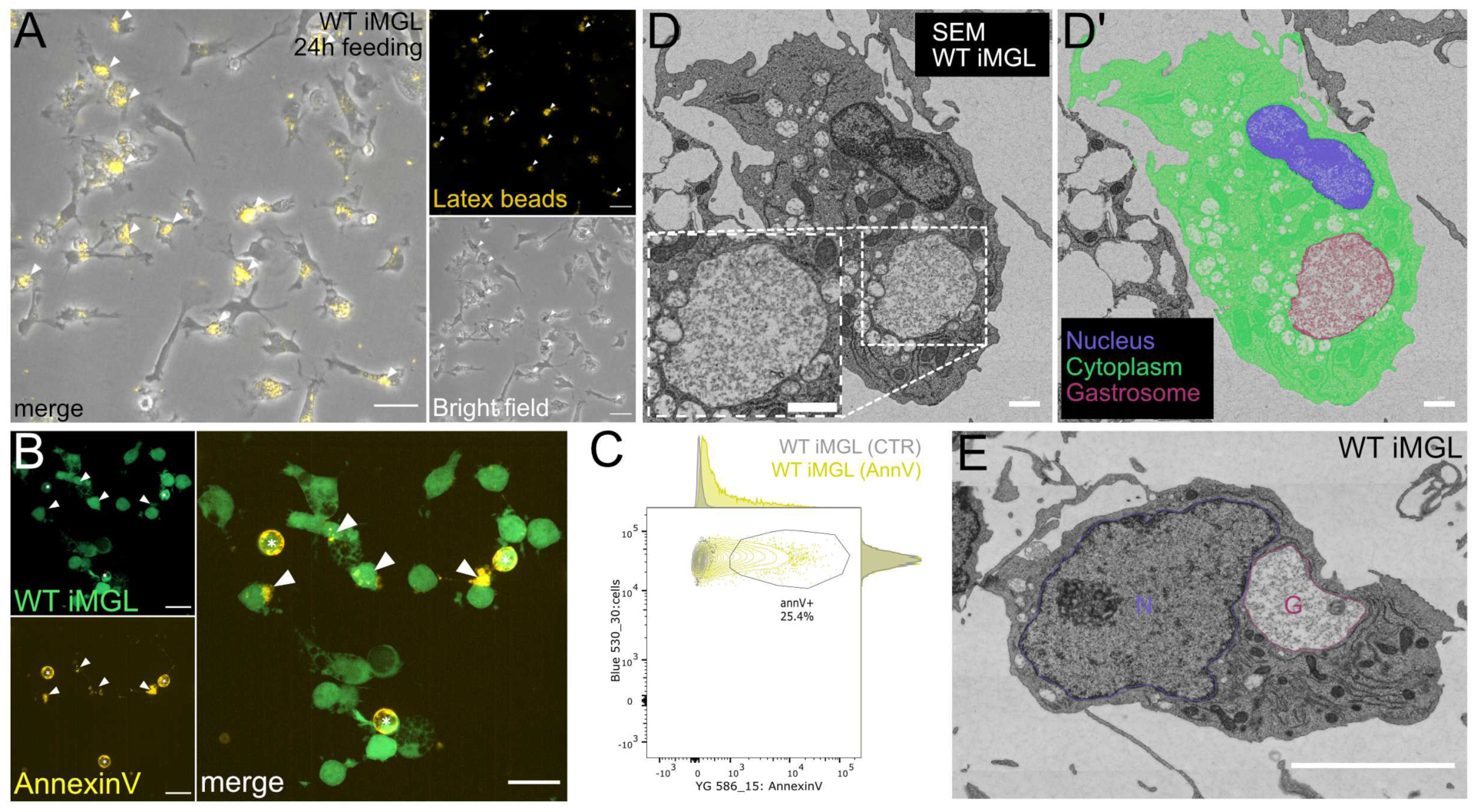
In iMGL the gastrosome is a collective phagocytic compartment characterized by distinct ultrastructure characteristics. **(A)** WT iMGL monoculture fed with 1 µm fluorescent latex beads and imaged after 24 h; scale bar 30 µm. **(B)** Light microscopy of WT iMGL monocultures stained with Annexin V to label apoptotic structures. Asterisks (*) indicate dead iMGL, while arrowheads mark Annexin V-positive debris within healthy, neighbouring iMGL; Scale bar: 20 µm. **(C)** Flow cytometry analysis of WT iMGL stained with Annexin V (yellow) compared to unstained control (gray). **(D)** Singe slice from an array SEM of WT iMGL in vitro (monoculture) with segmentation **(D’)** featuring the cytoplasm (green), nucleus (blue) and gastrosome (purple); scale bars 1 µm. **(E)** Electron micrograph single plane of WT iMGL in monoculture; N = nucleus, G = gastrosome; scale bar 5 µm.

**Supplementary Figure 4.**
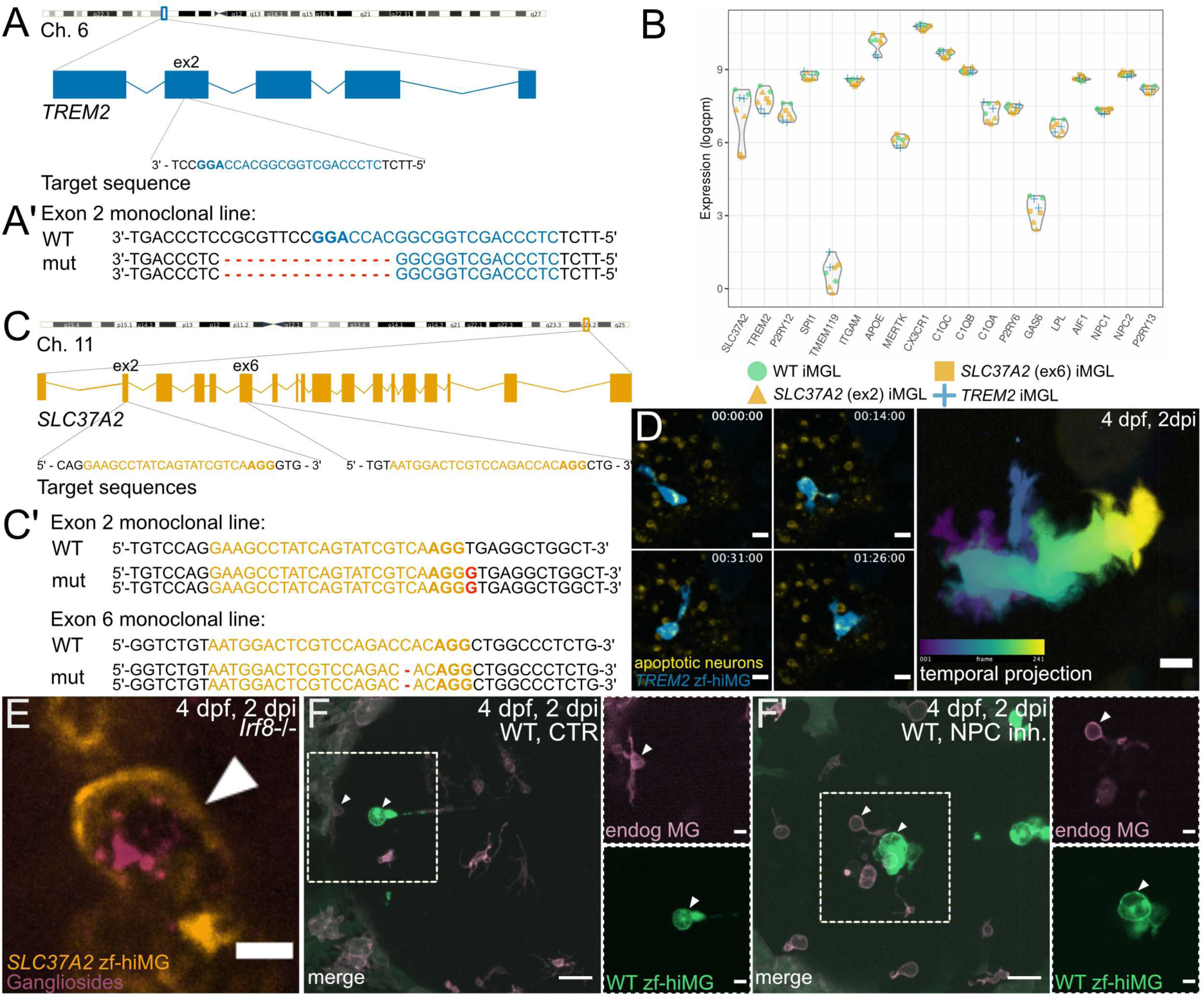
*In vitro* and *in vivo* characterization of perturbed human microglia-like cells. **(A)** Schematic of CRISPR-Cas9–mediated targeting strategy for TREM2 in GFP labelled human iPSCs, using a guide RNA designed to exon 2. **(A’)** Sequencing results of the TREM2 mutant clone (targeted exon). **(B)** Bulk RNA-sequencing analysis of WT, *SLC37A2* mutant and *TREM2* mutant iMGL, showing reduced *SL37A2* transcript levels in *SLC37A2* mutants compared to WT (ex2 log₂FC = –1.63, p = 0.000093, FDR = 0.056; not significant after correction; ex6 log₂FC = –3.37, p = 0.000000162, FDR = 0.00030, significant after correction), reduced *TREM2* transcript levels in *TREM2* mutants compared to WT (log₂FC = –1.37, p = 0.0000343, FDR = 0.0147, significant after correction) and preserved expression of canonical microglial markers for all genotypes. **(C)** Schematic of CRISPR-Cas9 targeting strategy for *SLC37A2* in GFP-labelled human iPSC, using a guide RNA designed to exon 2 and exon 6. **(C’)** Sequencing results of the two *SLC37A2* mutant clones (targeted exons). **(D)** Time lapse images of GFP-labelled *TREM2* zf-hiMG in *Irf8^st95^* embryos with neuronal apoptotic marker (Tg(nbt:dLexPR-LexOP:secA5-BFP); temporal projection of the same cell (right panel); dorsal view; 30 seconds time resolution; 4dpf, 2 dpi; scale bars 10 µm. **(E)** CtxB staining to detect GM1 gangliosides in GFP-labelled *SLC37A2* zf-hiMG xenotransplanted into *Irf8^st95^* zebrafish embryos; 4 dpf, 2 dpi; dorsal view; scale bar 10 µm. **(F-F’)** Morphological comparison of NPC1 inhibition **(F’)** versus control **(F)** in GFP-labelled WT zf-hiMG transplanted into WT embryos with microglia labelled with Tg(fms:Gal4; UAS:lyn-miRFP670); 4 dpf, 2 dpi; dorsal view; scale bars scale bars 30 µm (overview) and 10 µm (zoomed crop).

